# Atomic Force Microscopy Reveals Distinct Protofilament-scale Structural Dynamics in Depolymerizing Microtubule Arrays

**DOI:** 10.1101/2020.12.22.423986

**Authors:** Sithara S. Wijeratne, Michelle F. Marchan, Jason S. Tresback, Radhika Subramanian

## Abstract

The dynamic reorganization of microtubule-based cellular structures, such as the spindle and the axoneme, fundamentally depends on the dynamics of individual polymers within multi-microtubule arrays. A major class of enzymes, implicated in both the complete demolition and fine size control of microtubule-based arrays, are depolymerizing kinesins. How different depolymerases differently remodel microtubule arrays is poorly understood. A major technical challenge in addressing this question is that existing optical or electron-microscopy methods lack the spatial-temporal resolution to observe the dynamics of individual microtubules within larger arrays. Here we use Atomic Force Microscopy (AFM) to image depolymerizing arrays at single microtubule and protofilament resolution. We discover previously unseen modes of microtubule array destabilization by conserved depolymerases. We find that the kinesin-13 MCAK mediates asynchronous protofilament depolymerization and lattice-defect propagation, whereas the kinesin-8 Kip3p promotes synchronous protofilament depolymerization. Unexpectedly, MCAK can depolymerize the highly stable axonemal doublets, but Kip3p cannot. We propose that distinct protofilament-level activities underlie the functional dichotomy of depolymerases, resulting in either large-scale destabilization or length regulation of microtubule arrays. Our work establishes AFM as a powerful strategy to visualize microtubule dynamics within arrays and reveals how nanometer-scale substrate specificity leads to differential remodeling of micron-sized cytoskeletal structures.

**SIGNIFICANCE STATEMENT:** One cannot help but marvel at the precise organization of microtubule polymers in cellular structures such as the axoneme and the spindle. However, our understanding of the biochemical mechanisms that sculpt these arrays comes largely from in vitro experiments with a small number (one or two) of microtubules. This is somewhat akin to studying the architecture of multilane highways by studying one-lane streets. Here we directly visualize depolymerizing microtubule arrays at individual microtubule and protofilament resolution using Atomic Force Microscopy. Our results reveal differences in microtubule depolymerase activity and provide insights into how these differences in enzymatic activity on the nanometer scale can result in the differential remodeling of multi-microtubule arrays on the micron-length scale.

## MAIN TEXT

The dynamic formation and dismantling of protein arrays underlie a broad range of cellular functions in both prokaryotes and eukaryotes. A prototypical example of dynamic polymeric protein structures are micron-sized arrays of microtubules, which assemble into essential cellular machines and tracks such as the mitotic spindle in dividing cells, axonal arrays in neurons and axonemes in cilia and flagella. Microtubules themselves are complex cylindrical macromolecular assembly of, most commonly, 13-15 protofilaments that are composed of repeating α,β-tubulin heterodimers. The intrinsic dynamic instability of microtubules and its regulation by a host of different Microtubule Associated Proteins (MAPs) are critical for the assembly and disassembly of microtubule arrays (1). How nanometer-scale dynamics of protofilaments (~4 nm) and microtubules (~25 nm) result in the organization and remodeling of micron-sized multi-microtubule arrays remains poorly understood.

In vitro reconstitution and visualization by optical microscopy have provided tremendous insights into microtubule dynamic instability and its regulation by MAPs. However, these studies have been limited to single or pairs of microtubules as light microscopy does not have the resolution to identify individual microtubules within a complex array of multiple microtubules. In addition, it is challenging to image individual protofilaments within each microtubule by this method. Structural intermediates of microtubule-remodeling reactions have been inferred from electron microscopy studies, but the single snapshot nature of the technique lacks temporal resolution to follow reaction dynamics in real time. To address these technical limitations and offer insights into microtubule array remodeling at single microtubule and protofilament resolution in real time, we employed Atomic Force Microscopy (AFM) imaging (2–8). This technique allowed the direct visualization of microtubule depolymerization in two different arrays, antiparallel microtubule bundles that are found in the mitotic spindle and doublet microtubule arrays that form axonemes in cilia and flagella (9–11).

A critical reaction that governs the size and stability of microtubule arrays is microtubule depolymerization, which is catalyzed by a class of enzymes known as microtubule depolymerases. This reaction is required for rapid large-scale reorganization of the cytoplasm. For example, the mitotic spindle is built and disassembled every time a cell divides and the cilium is constructed and deconstructed each cell cycle (12–14). In addition to large scale reorganization of microtubule networks and arrays, microtubule dynamics and its regulation are important for fine tuning the size of microtubule arrays (15–18). A fundamental conundrum is how the same reaction, the removal of tubulin from microtubules, results in different outcomes ranging from large-scale remodeling or fine-length regulation of microtubule arrays.

Prototypical depolymerases are members of the two major family of kinesins, the vertebrate kinesin-13 family, such as MCAK, and the budding yeast kinesin-8 family, such as Kip3p (19–21). While the non-motile MCAK and processive Kip3p proteins have different mechanisms for arriving at the microtubule ends, enzymatically MCAK and Kip3p are both catastrophe factors and catalyze the removal of tubulin from microtubule ends (22, 23) (24, 25). Structural studies suggests that at the microtubule end, both enzymes recognize the curved conformation of tubulin in a similar manner (26–28). Despite these similarities, these proteins differently regulate dynamic instability such that Kip3p limits the distribution of maximum microtubule lengths whereas MCAK promotes rapid filament shortening (29–31). These differences are reflected in their distinct functions; kinesin-8 proteins are largely involved in length control of structures such as the spindle and the cilium, while kinesin-13s are additionally implicated in large-scale cytoskeleton remodeling such as the depolymerization of interphase microtubules during entry into mitosis and suppression of cilium biogenesis (32–36). However, what underlies the differences in activity of these prototypical kinesin-family depolymerases and how differences in depolymerase activity at the single microtubule level translate to distinct remodeling of complex arrays remain unknown.

The AFM imaging reported here reveals the structural dynamics that underlie microtubule array destabilization and provide a framework for linking the action of enzymes on the nanometer-scale protofilaments to the remodeling of micron-sized arrays. The study sheds light on the long-standing question of how different depolymerases are tuned for distinct cellular activities, such as rapid remodeling or length control of microtubule arrays. Our findings highlight differences in enzyme activity on the protofilament scale as a critical parameter that governs the fate of microtubules within complex structures, thereby dictating how such arrays are remodeled.

### Imaging the depolymerization of PRC1-crosslinked microtubule arrays by Atomic Force Microscopy

As a first step towards imaging the depolymerization of microtubule arrays by MCAK, we reconstituted microtubule bundles using the antiparallel crosslinking protein PRC1 (Protein Regulator of Cytokinesis-1) on a mica surface (Fig. 1A) (see Methods and SI Appendix, Methods and Materials, Extended Methods). AFM images of the bundles reveal that PRC1 crosslinking results in 2D microtubule arrays (Fig. 1B-C). This is advantageous as every microtubule in the bundle can be clearly distinguished visually (see Methods and SI Appendix, Methods and Materials, Extended Methods). This is reflected in a height plot as a series of peaks with an average height of ~30 nm (Fig. 1D). The 3D rendition of the images, obtained from the surface topography data of Fig. 1B, shows dense linkages connecting the overlapping microtubules in an array, consistent with the cooperative binding of PRC1 to overlapping microtubules (Fig. 1C) (37, 38).

**Fig. 1.**
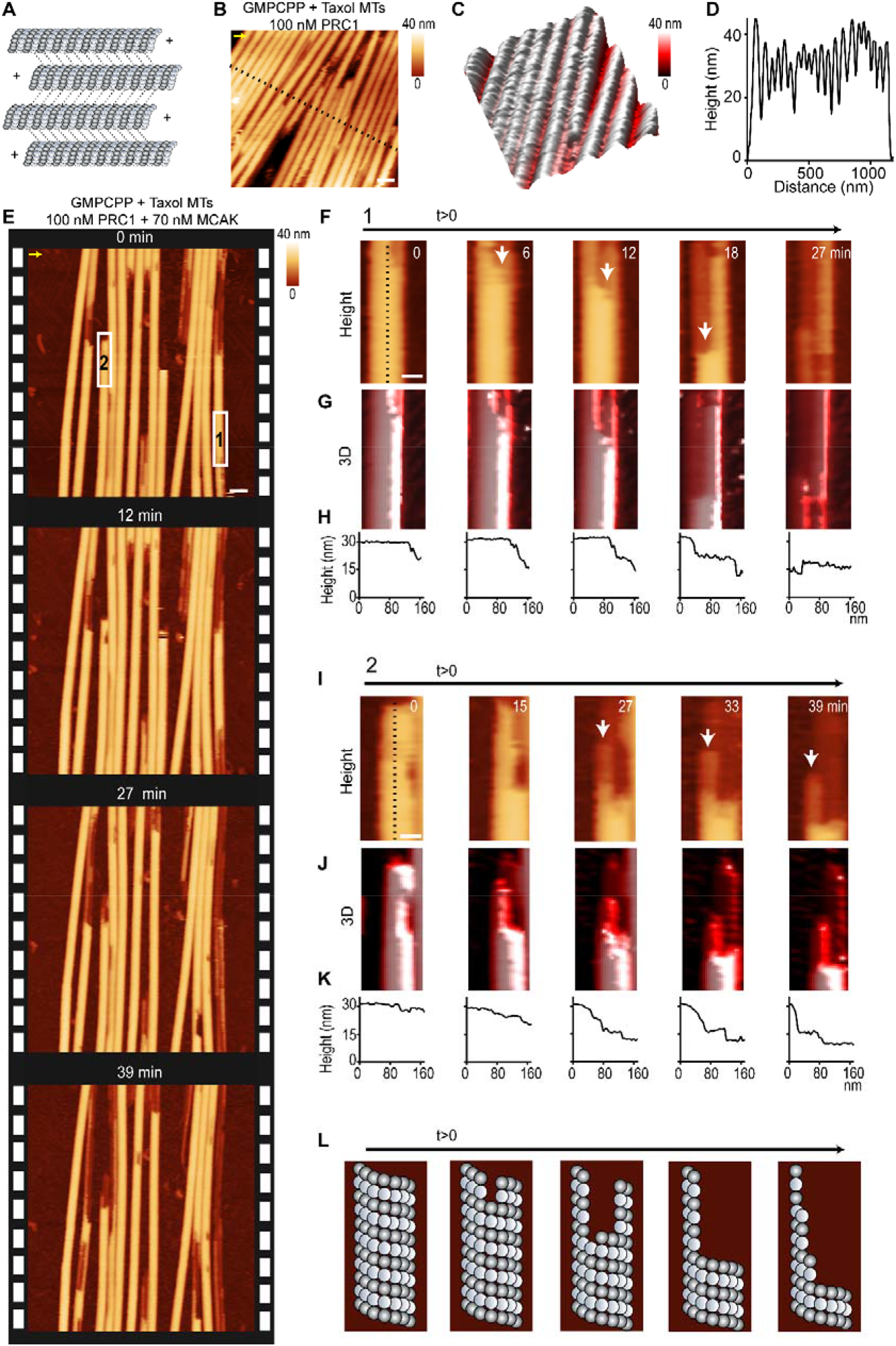
MCAK depolymerizes microtubule protofilaments asynchronously. (A). Schematic of an antiparallel microtubule array crosslinked by PRC1 (dotted lines). (B). AFM image of a microtubule bundle crosslinked by 100 nM PRC1. Each microtubule within the flat 2D array can be clearly distinguished. The x-y scale bar is 70 nm. The z-scale is 0 to 40 nm (dark to light brown). The AFM image is colored according to height from the surface. (C). The 3D rendition of a zoomed in region from B. (D). The corresponding height profile from dotted line in B. (E). Successive AFM images show depolymerization of individual microtubules within a PRC1-crosslinked bundle by MCAK from 0-39 min. The image at 0 min represents the first frame taken after adding MCAK (GMPCPP + Taxol microtubules, PRC1:100 nM; MCAK: 70 nM). The x-y scale bar is 100 nm. The z-scale is 0 to 40 nm (dark to light brown). (F)&(I). Two examples of depolymerization from the experiment in E showing stripe-like appearance of depolymerizing protofilaments (boxes 1-2). (G)&(J). The 3D rendition of the AFM time-lapse images from F and I. (H)&(K). Corresponding height profiles from the white dotted line in panels F and I show that the stripiness results from protofilaments at different heights relative to the surface. (L). Schematic of a microtubule undergoing asynchronous protofilament-level depolymerization. The scanning rate is 4 min/frame in B and ~3 min/frame in E with 256 x 256 pixels. The arrow on panels B) and E) indicates the scanning direction in the fast axis. See also SI Appendix, Figs. S1-3.

We used two main criteria to set the imaging conditions for investigating microtubule dynamics within PRC1-crosslinked bundles (3-4 mins/frame with 256 x 256 pixels): (1) the spatial and temporal resolution is suitable for imaging the entire array as well as individual microtubules within the bundle, and (2) no sample damage is visible in the time frame of the experiment (15-30 mins). Experiments were performed with both GMPCPP- and GMPCPP + taxol-stabilized microtubules. After locating a microtubule bundle by AFM, MCAK was added into the sample chamber at the indicated concentrations and time lapse image series was acquired (*note:* solution concentrations are reported throughout the manuscript; local concentrations on the mica surface may be different). The rate of microtubule depolymerization by MCAK in the absence of PRC1 is similar to the rates observed by TIRF (SI Appendix, Fig. S1A-C), which shows that the activity of the enzyme is intact in this assay (SI Appendix, Fig. S1D-E) (24, 29, 39). The time-lapse AFM imaging of PRC1-crosslinked bundles reveals that depolymerization of individual microtubules within an array by MCAK can be visualized in real time (Fig. 1E, SI Appendix, Fig. S2, Videos 1-2). Depolymerization begins shortly after MCAK addition, as we observed depolymerization of microtubules within the arrays in the first image taken after MCAK addition. This is visualized in sections of microtubule that lack complete tubules (Fig. 1E, time=0 min). In control experiments without MCAK, PRC1-crosslinked microtubules remain stable and do not depolymerize significantly during the experiment (SI Appendix, Fig. S3).

These observations reveal AFM imaging as a powerful method to examine dynamic changes in a multi-microtubule array with nanometer-scale spatial resolution, thus offering a view of structural changes that are extremely difficult to detect by other imaging methods (40).

### MCAK depolymerizes microtubule protofilaments asynchronously and propagates defects within crosslinked bundles

Close examination of the time lapse AFM images revealed two striking features of the reaction intermediates. First, the loss of microtubules was observed to be associated with the appearance of stripe-like features in the AFM images, which correspond to protofilaments. For example, in the magnified time-lapse montages, partial and bidirectional depolymerization is observed at the microtubule ends (Fig. 1F-K, SI Appendix, Fig. S2A). The stripes arise due to protofilaments at different heights from the surface, which are resolved because the high resolution of AFM in the axis perpendicular to the mica surface (see SI Appendix, Extended Methods). These observations show that in the presence of MCAK, the depolymerization of protofilaments within a microtubule is asynchronous (Fig. 1F, 1I, SI Appendix, Fig. S2A). The stripiness is observed on non-crosslinked as well as crosslinked microtubules with one or two neighbors. These observations suggest that microtubule protofilaments depolymerize at different rates in the presence of MCAK, resulting in an asynchronous loss of protofilaments from the ends (Fig. 1L). To ensure that the features observed do not arise from the frequency of AFM scanning, we increased the time interval between scans to 5-10 mins after MCAK addition and found that the features were unaltered and independent of time interval (SI Appendix, Fig. S2D-E). Similar results were observed with microtubules stabilized by either GMPCPP or GMPCPP + taxol (Fig. 1F, SI Appendix, Fig. S2). EM studies have described curved ram horn shape of protofilaments during depolymerization. In the AFM experiments, these transient structures can only be captured if they lie in the plane of the mica surface as shown in SI Appendix, Fig. S2B.

Second, in addition to depolymerization from the ends, we observed that breaks can also appear in the middle of microtubule arrays. These are likely to be defects formed by the loss of tubulin from the microtubule lattice. We find that defects can be propagated by MCAK, again accompanied by the protofilament-associated stripiness (Fig. 2A-C, SI Appendix, Fig. S2B). Defects propagate both along the diameter and length of microtubules, with different rates (Fig. 2D-E), suggesting that destabilization of inter-protofilament interactions is slower than depolymerization along the length of an individual protofilaments in the presence of MCAK. The observed depolymerization rates in the two opposite directions along the microtubule length are consistent with the unequal rates of plus and minus end microtubule depolymerization by MCAK (Fig. 2D) (23). More defects were observed in our taxol stabilized samples compared to GMPCPP, suggesting that polymerization conditions contribute to defects (41). Defect propagation was also observed in Total Internal Reflection Fluorescence (TIRF) microscopy-based assays with GMPCPP microtubules in the presence of MCAK (300-500 nM) (SI Appendix, Fig. S1A-C). Thus, while the number of defects observed in AFM may be a combination of pre-existing lattice defects and any additional ones induced by the AFM tip, the TIRF and AFM data show that MCAK can propagate these defects to induce destabilization of arrays.

**Fig. 2.**
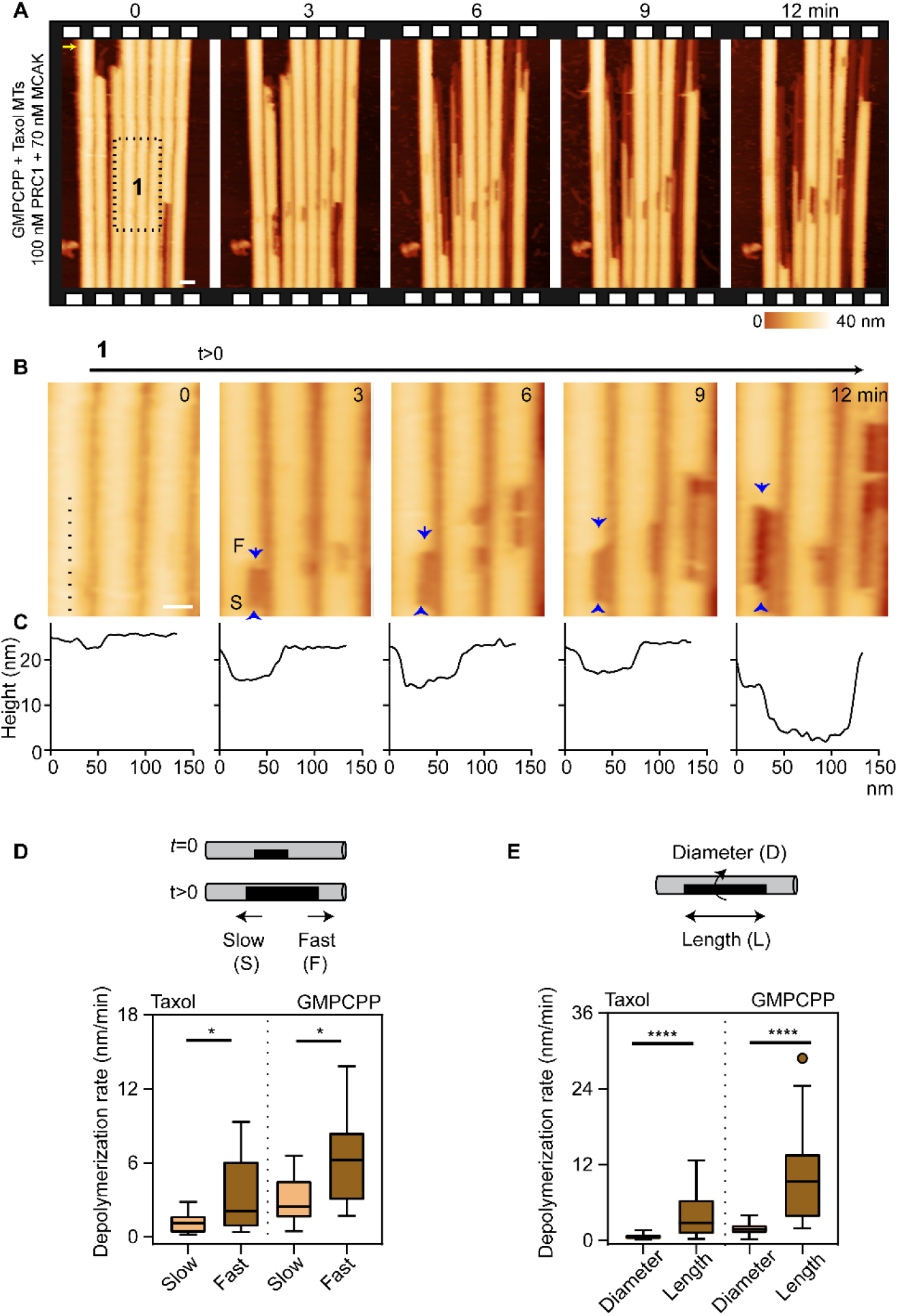
MCAK propagates defects within crosslinked bundles. (A). Successive AFM images showing depolymerization of individual microtubules within a microtubule bundle by MCAK at 0-12 min. The image at 0 min represents the first frame taken after adding MCAK (GMPCPP + Taxol microtubules, PRC1:100 nM; MCAK: 70 nM). The x-y scale bar is 50 nm. (B). Successive time-lapse montages of a zoomed-in region of a section of microtubule from the experiment in A (box 1) showing defect propagation on three microtubules within the array at the indicated times (arrows). ‘S’ indicates the edge with slow depolymerization, and ‘F’ indicates the edge with fast depolymerization. The x-y scale bar is 30 nm. (C). The corresponding average height profiles along the length of the microtubule from the dotted line in B. (D). Box plots of the depolymerization rates of ‘slow (S)’ and ‘fast (F)’ events from defect propagation with MCAK (Median: PRC1: 100 nM; MCAK: 70 nM; GMPCPP + Taxol microtubules: slow=1 nm/min, n=19, fast=2 nm/min, n=17; GMPCPP microtubules: slow= 3 nm/min, n=18, fast=6 nm/min, n=18). (E). Box plots of the depolymerization rates of ‘diameter (D)’ and ‘length (L)’ events from defect propagation with MCAK (Median: PRC1: 100 nM; MCAK: 70 nM; GMPCPP + Taxol microtubules: diameter=0.6 nm/min, n=20, length=3 nm/min, n=20; GMPCPP microtubules: diameter= 2 nm/min, n=22, length=10 nm/min, n=22). The scanning rate is ~3 min/frame with 256 x 256 pixels. The arrow on panel A) indicates the scanning direction in the fast axis. For D-E, the box plots show the median, the inner quartiles, and maximum and minimum values. See also SI Appendix, Fig. S2. Statistical calculations used an unpaired t-test with Kolmogorov-Smirnov correction for non-Gaussian distribution. * indicates a P-value of <0.05. **** indicates a P-value of <0.0001.

How does PRC1-mediated bundling of microtubules affect array destabilization by MCAK? To address this, we quantitatively examined the effect of neighboring microtubules in bundles on the depolymerization reaction. Quantitative measurement of the depolymerization rates of microtubules that have zero, one or two neighbors (see SI Appendix, Methods and Materials) showed that bundling has a protective effect on microtubules against MCAK-mediated depolymerization (SI Appendix, Fig. S1E). We find that the depolymerization rates of microtubules with two neighbors are 3-fold lower than microtubules with zero or one neighbor. Second, microtubule depolymerization rates also depend on PRC1 concentration (SI Appendix, Fig. S1E). For instance, the depolymerization rate decreased when the solution concentration of PRC1 was increased by 10-fold from 10 nM to 100 nM. This reduction could be alleviated by increasing the MCAK concentration by ~10-fold. Under all conditions, the presence of two neighboring microtubules has a significant effect on the protection of bundles. This is likely to arise from the highly dense pattern of PRC1 occupancy in overlap regions. Consequently, microtubules with two neighbors, one on either side, are likely to have the least number of exposed plus/minus protofilament ends (hereafter referred to as “exposed protofilament ends”).

The features observed during AFM imaging of microtubule array depolymerization by MCAK suggest that a single or few protofilaments with exposed ends can be an effective substrate for MCAK, and that these protofilaments are asynchronously removed by the enzyme. Altogether, we demonstrate that microtubule depolymerization can be visualized by AFM in real time at the single microtubule and protofilament resolution within arrays. By using this imaging modality, we provide the first view of protofilament level depolymerization by a microtubule depolymerase, nearly two decades after such possibility was hypothesized (42, 43). Therefore, we formally show that protofilament-level depolymerization occurs on microtubules within a larger array in the presence of MCAK. The observed asynchronous loss of protofilaments and defect propagation suggests that MCAK can use an exposed protofilament ends as a substrate as would be advantageous for large-scale destabilization of the microtubule arrays.

### Structural dynamics of microtubule depolymerization by Kip3p are distinct from MCAK

To investigate whether different depolymerases exhibit distinct structural dynamics, we examined the depolymerization of PRC1-crosslinked microtubule bundles by Kip3p. We mainly focused on GMPCPP stabilized microtubules since depolymerization of doubly-stabilized GMPCPP-taxol microtubules by Kip3p is extremely slow (23, 24). We first examined Kip3p activity on single microtubules in the absence of PRC1 (SI Appendix, Fig. S4A). The depolymerization rate increases with Kip3p concentration (SI Appendix, Fig. S4B) and are similar to the rates observed by TIRF microscopy (SI Appendix, Fig. S4C-D). Depolymerization is primarily seen at one end of the microtubules although we also observe some slow minus-end depolymerization (SI Appendix, Fig. S4B inset; 20 nm/min on the plus end compared to 5 nm/min on the minus end at 1 nM solution concentration of Kip3p). Low levels of minus depolymerization are also observed in TIRF assays at concentrations > 50 nM (SI Appendix, Fig. S4C-D). Enhanced local concentration of Kip3p (on the mica substrate or on the cantilever tip) may contribute to Kip3p-induced depolymerization activity from both ends of the microtubules as has also been observed with a non-motile Kip3p mutant (26). Control experiments confirm that the depolymerization in this experiment is enzyme mediated (SI Appendix, Fig. S4E). We next examined Kip3p activity on PRC1-crosslinked microtubules. We selected a field where the ends of several microtubules within the PRC1 bundle were in view, added Kip3p, and collected time-lapse AFM images (Fig. 3A, SI Appendix, Fig. S5A-B, Video 3). We observed that the structural features of depolymerization of PRC1-bound crosslinked and individual microtubules by Kip3p were distinct from MCAK. First, in contrast to MCAK, no stripes were observed at the ends of depolymerizing microtubules and the protofilaments were lost in unison, leaving only one or few protofilament remnants (< 5 nm in height) that were adhered on the mica surface (Fig. 3B-D). This is reflected in the shifting of the entire edge of the corresponding height profiles with depolymerization without features of intermediate heights. Second, unlike with MCAK, we rarely observed defect propagation events using the same batch of microtubules. In the rare instances where we saw depolymerization from the middle of a doubly stabilized GMPCPP-taxol microtubule, we again observed no stripes suggesting that defects are propagated by Kip3p in unison, i.e. synchronously (SI Appendix, Fig. S5B). Third, in contrast to MCAK, we did not see a significant effect of PRC1 concentration or neighbors on depolymerization rate (SI Appendix, Fig. S5C). Control experiments in the presence of ATP alone, confirm that the observed depolymerization of PRC1-crossslinked bundles in the presence of Kip3 is specific to the presence of depolymerases in the assay (SI Appendix, Fig. S3).

**Fig. 3.**
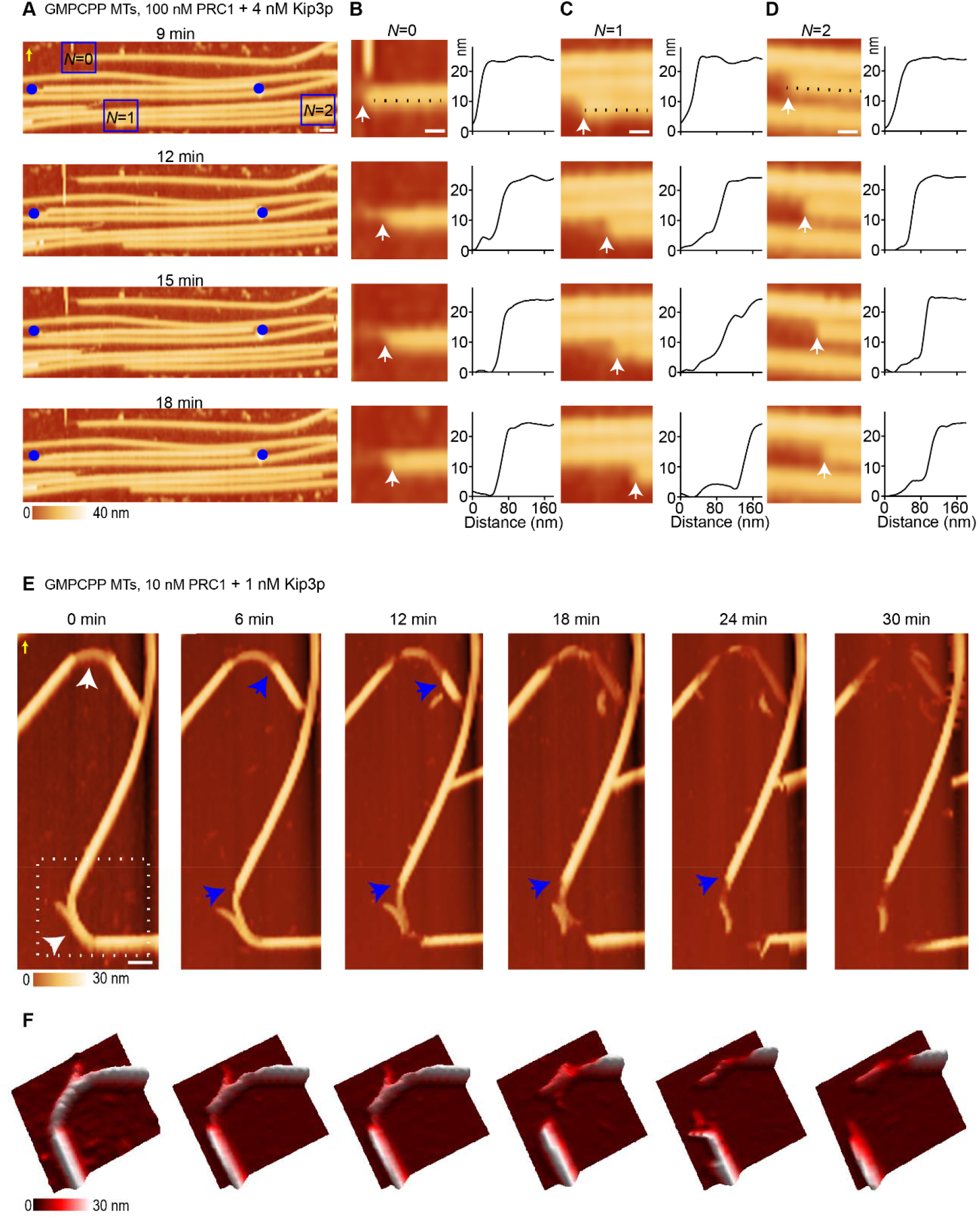
Structural dynamics of microtubule depolymerization by Kip3p are distinct from MCAK. (A). Successive AFM time lapse images of a PRC1-crosslinked microtubule bundle in the presence of Kip3p at the indicated times (GMPCPP microtubules, PRC1:100 nM; Kip3p: 4 nM). The blue circles are fiduciary marks which show that the microtubules are depolymerizing and not sliding. The x-y scale bar is 100 nm. (B)-(D). Zoomed-in regions from the experiment in A (boxes) showing bluntness at the depolymerizing microtubule end for N=0 (B), N=1 (C), N=2 (D) events. The height profiles corresponding to the dotted lines show that protofilaments at the ends of the microtubules are lost synchronously (white arrows). The x-y scale bar is 40 nm. For A-D, the z-scale is 0 to 40 nm (dark to light brown). (E). Successive AFM time-lapse images show the destabilization of two highly curved regions (white arrows at 0 min) at the indicated times. Over time, the microtubule starts to depolymerize faster from one end with synchronous loss of protofilaments (blue arrows) (GMPCPP microtubules, PRC1:10 nM; Kip3p: 1 nM). The x-y scale bar is 100 nm. The z-scale is 0 to 30 nm (dark to light brown). (F). The 3D rendition shows a magnified view of this depolymerization activity (dotted box at 0 min in E). The scanning rate is ~3 min/frame with 256 x 256 pixels. The arrow on panels A) and E) indicates the scanning direction in the fast axis. See also SI Appendix, Figs. S4-S6.

In addition, we observed that Kip3p destabilizes highly curved microtubules, consistent with the reported accumulation of Kip3 at these sites (26) (Fig. 3E-F, SI Appendix, Fig. S6A). Destabilization of the curved region was neither observed in the presence of the motile conventional kinesin (K401) or ATP alone (SI Appendix, Fig. S6B-C). We used time-resolved AFM to follow the structural intermediates of the process by which the curved segment disintegrates in real time in the presence of Kip3p (Fig. 3E-F, SI Appendix, Video 4) (26). As seen in the zoomed 3D view, the curved section destabilized and depolymerized faster in one direction (Fig. 3F). Again, we observed that depolymerization of the entire microtubule occurs when most of the protofilaments in the curved region are lost. Overall, the features of microtubule depolymerization, such as synchronous depolymerization of protofilaments, lack of defect propagation, and accelerated breaking at curved microtubule segments by Kip3p in the absence of PRC1 (SI Appendix, Fig. S4A, S6A), are similar to that observed in experiments with PRC1 (Fig. 3A-F, SI Appendix, Fig. S5A-B). Together these data suggest that unlike MCAK, Kip3p depolymerizes protofilament ends synchronously. This is likely due to the accumulation of Kip3p at microtubule ends and a preference to stall at ends, rather than at sections of partially exposed protofilament ends such as those in defects.

Altogether, this suggests that the two depolymerases, Kip3p and MCAK, exhibit distinct preferences in terms of substrate specificity at microtubule ends and defects at the protofilament level, and in the context of an array of bundled microtubules. These findings shed light on how the two depolymerases may be tuned for distinct functions. While the properties of MCAK make it well suited for remodeling of arrays and depolymerization at defects, Kip3p activity seems better aligned with a role of a length regulator at microtubule ends.

### Visualizing the depolymerization of doublet-microtubules using AFM

As a step towards examining how other microtubule arrays are remodeled by depolymerases, we focused on the axoneme, an array of nine outer doublet and two central singlet microtubules that forms the backbone of the cilia and flagella (Fig. 4A). The doublets are composed of the A-tubule, which contains 13 protofilaments, and the B-tubule, which is an incomplete microtubule containing ten protofilaments (44, 45). At the distal cilium tip, the doublets transition into an array composed of singlets (44). Axoneme are one of the most stable arrays of microtubules in cells and dissociating them into soluble tubulin requires fairly harsh treatments like sonication and detergent, or specific ionic conditions (46, 47). Depolymerases of the kinesin-13 and kinesin-8 families are proposed to act on axonemes to control cilium length and stability (32–34). However, the activity of depolymerases on doublet microtubules has not been visualized, and it is unknown if and how these enzymes depolymerize doublets and impact axoneme stability.

**Fig. 4.**
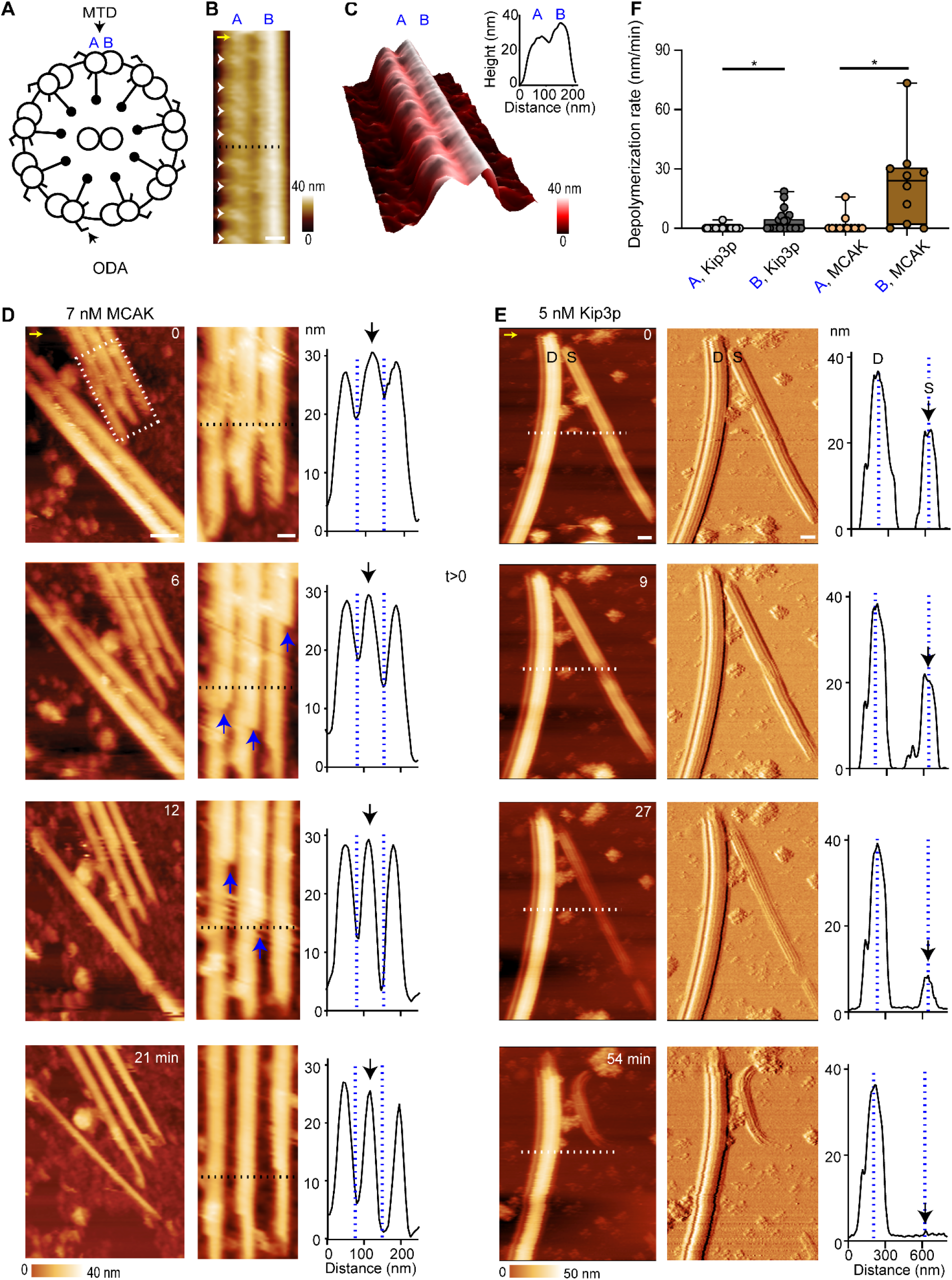
Visualizing the activity of MCAK and Kip3p on doublet-microtubules using AFM. (A). Schematic of the axoneme structure. Axonemes consists of 9 outer microtubule doublets (MTD). Each doublet contains an A and B tubule. Outer dynein arms (ODA) present on the A tubule form a repeating pattern. (B). AFM height image of an MTD. The A and the B tubule in an MTD were distinguished by the height of the tubules in the joined doublet (25 and 35 nm) and the periodic striations, which are separated by 30 nm (arrows). The x-y scale bar is 50 nm. (C). The 3D representation of B and the corresponding height profile of dotted line in B. (D). Successive AFM images show a 2D microtubule doublet sheet with MCAK at the indicated times. The zoomed in region (boxed region) shows the depolymerization activity of alternate tubule in an array (arrows) and its corresponding height profiles over time (dotted line). The height profiles show the deepening of the minima and changing of the asymmetric peak into a single sharp peak (dotted line, arrow) (-TED sample, MCAK: 7 nM). The x-y scale bar of the zoomed-out and zoomed-in image is 80 nm and 40 nm, respectively. The z-scale is 0 to 40 nm (dark to light brown). (E). Successive AFM height and amplitude images show a microtubule doublet (D) and a singlet (S) in the presence Kip3p at the indicated times. The corresponding height profiles from dotted line show that the height of the doublet doesn’t change over time, but the height of the singlet reduces with time (dotted lines) (-TED sample, Kip3p: 5 nM). The x-y scale bar is 100 nm. The z-scale is 0 to 50 nm (dark to light brown). (F). Box plots of the depolymerization rates of A and B tubules in the -TED sample with MCAK and Kip3p. MCAK: A rate=2 nm/min, n=12; B rate=23 nm/min, n=10; Kip3p: A rate=0.2 nm/min, n=22; B rate=3 nm/min, n=22. Data were pooled from experiments with protein concentrations less than 10 nM. Statistical calculations used an unpaired t-test with Kolmogorov-Smirnov correction for non-Gaussian distribution. * indicates a P-value of <0.05. The scanning rate is ~4 min/frame in B and ~3 min/frame in E-F with 256 x 256 pixels. The arrow on panels B), D) and E) indicates the scanning direction in the fast axis. For F), the box plot shows the median, the inner quartiles, and maximum and minimum values. See also SI Appendix, Figs. S7 and S8.

We purified axonemes from *Lytechinus pictus* sea urchin sperm. We first focused on individual doublets present in this sample. In high spatial resolution mode, the AFM images clearly show two joined tubules, with heights of 25 and 35 nm respectively (Fig. 4B-C). Another feature of these microtubule doublets were the periodic repeats separated by ~30 nm, which are likely to be the outer dynein arms on the A tubule (Fig. 4B-C). Both of these structural features of microtubules are consistent with cryo-EM structures of the axoneme and previous AFM images of doublet microtubules (48, 49). First, we examined whether depolymerases act on doublets microtubules by performing AFM experiments with 2D doublet sheets from partially dissociated axonemes, composed of two or more doublets linked together (SI Appendix, Fig. S7A-D). In the presence of MCAK, we observed that one of the tubules depolymerizes from both ends first, followed by the depolymerization of the second tubule (Fig. 4D, SI Appendix, Figs. S7E-F and Video 5). This can be visualized in the height plots as a deepening of the minima between adjacent doublets (dotted lines) and the conversion of an asymmetric broad peak to a single sharp peak (indicated by black arrow) (Fig. 4D). The AFM time-lapse data of a microtubule doublet and a singlet in the same field of view in the presence of Kip3p revealed that the doublet depolymerizes very slowly (rate= 4 nm/min), but the single microtubule is depolymerized on the same time scale (rate= 26 nm/min) (Fig. 4E and SI Appendix, Video 6). This is consistent with previous fluorescent studies with the kinesin-8 protein Kif19A, where single microtubules nucleated from an axoneme were depolymerized, but the axoneme itself appears intact (32). Within the doublet, the average depolymerization rate measured for the A- and B-tubule in the presence of Kip3p was close to zero (0.2-3 nm/min) (Fig. 4F), while we estimated that MCAK leads to faster B-tubule depolymerization than the A-tubule (Fig. 4F; A rate= 2 nm/min; B rate= 23 nm/min). Control experiments with the doublets without enzyme showed no significant depolymerization over the same time scale (SI Appendix, Fig. S7G). Taken together, these data suggest clear differences in how MCAK and Kip3p process microtubule doublets.

Axonemes are heavily decorated with several proteins. Prior reports have shown that treatment with a solution of Tris-EDTA-DTT (TED) can dissociate a subset of axonemal proteins, particularly the dyneins (50). Our AFM imaging revealed that these TED-treated doublets have a smoother surface in comparison to untreated doublets (SI Appendix, Fig. S8A-B). We examined if TED treatment alters the depolymerization of doublets. Similar to the doublet samples without TED treatment (Fig. 4), we observed bidirectional depolymerization of one tubule of an isolated doublet with MCAK (SI Appendix, Fig. S8C). TED treatment resulted in faster depolymerization of both tubules (compared to non-TED with MCAK) with the B-tubule being lost at a higher rate than the A-tubule (SI Appendix, Fig. S8D; A rate= 16 nm/min; B rate= 38 nm/min). In the presence of Kip3p, TED treatment also permitted slow depolymerization of the B-tubule, but the A tubule remained intact (SI Appendix, Fig. S8E-F; A rate= 2 nm/min; B rate= 10 nm/min). To ensure that depolymerization is not arising from frequent AFM scanning, we have imaged the doublets every 10 minutes upon adding a depolymerase (SI Appendix, Fig. S8G). The results show that the depolymerization activity of the doublet with MCAK is independent of AFM scanning. In addition, in control experiments, without the enzyme, TED-treated doublets showed no significant depolymerization (SI Appendix, Fig. S8H).

Taken together, our results show for the first time that doublet microtubules can be enzymatically depolymerized with MCAK, and the rate of depolymerization of the B-tubule in a doublet is faster than the A-tubule. In contrast, doublet microtubules are poor substrates for Kip3p even when the doublet is stripped of associated proteins. Thus, proteins that have Kip3p-like properties may be selectively functional at the distal cilium tip, where the axoneme is composed mostly of singlet microtubule, to fine-tune cilium length. In contrast, the properties of depolymerases like MCAK make them better suited to depolymerize both singlets and doublets for processes such as cilia disassembly or inhibition of ciliogenesis.

### Destabilization of axonemal structures by MCAK

Our observations with the doublet microtubule depolymerization raise the question of how preferential depolymerization of B-tubule by MCAK impacts stability of the entire axoneme. We first imaged intact axonemes adsorbed onto the mica surface. Axonemes were ~200 nm in height, which is consistent with the diameter of the axoneme from cryo-EM measurements (Fig. 5A-D and SI Appendix, Fig. S9A-F) (48). The AFM amplitude image showed longitudinal striations, which likely arise from the nine outer doublets that are around ~20-30 nm apart (Fig. 5B and SI Appendix, Fig. S9B). In AFM imaging experiments with ATP alone, we observed no substantial change in the overall size and integrity of the axoneme over time (Fig. 5E).

**Fig. 5.**
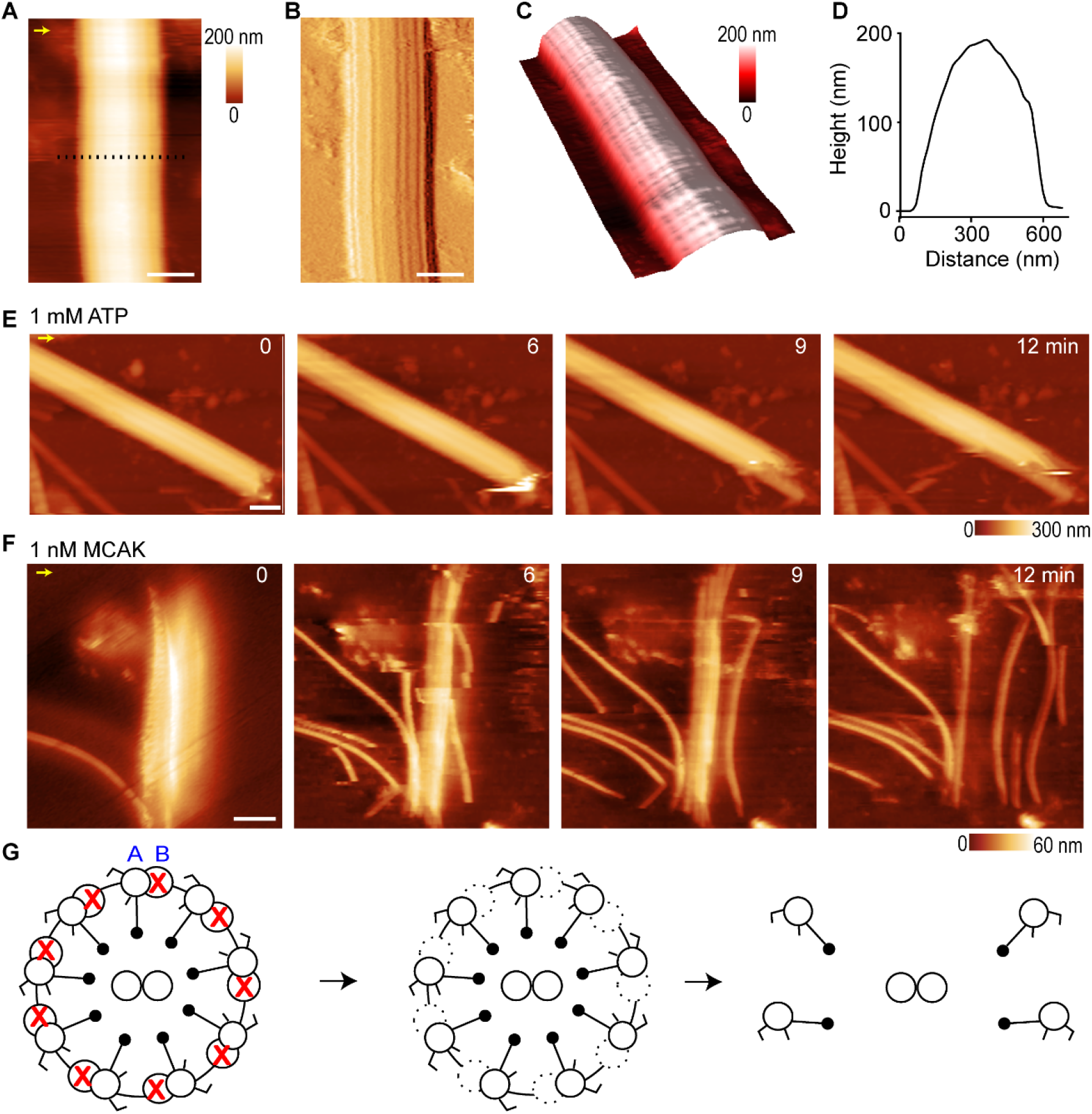
Destabilization of axonemal structures by depolymerases. (A)-(C). AFM height (A), amplitude (B) and 3D (C) images of an intact axoneme from Lytechinus pictus sea urchin sperm. The AFM amplitude (B) and the 3D (C) images show longitudinal striations, likely from the nine outer doublets, which are around ~20-30 nm apart. The x-y scale bar is 300 nm. The z-scale is 0–200 nm (dark to light brown). (D). The height profile of the selected dotted line from height image in A is ~200 nm. (E). Successive AFM images of an intact axoneme with 1 mM ATP at the indicated times. With ATP alone, no significant change was observed in the axoneme structure. The x-y scale bar is 300 nm. The z-scale is 0–300 nm (dark to light brown). (F). Successive AFM images of an axoneme in the presence of MCAK (1 nM) at the indicated times. At t=0, the axoneme has been partially frayed. The x-y scale bar is 500 nm. The z-scale is 0–90 nm (dark to light brown). (G). Schematic of proposed intermediate that results in unfurling of the axoneme with MCAK. The scanning rate in A-C is ~4 min/frame in B and ~3 min/frame in E-F with 256 x 256 pixels. The arrow on panels A), E) and F) indicates the scanning direction in the fast axis. See also SI Appendix, Fig. S9.

Next, we selected an axoneme, added depolymerase (t = 0), and monitored the changes in the axoneme structure with time. With the addition of MCAK, we observed a rapid loss of microtubules from the axoneme. For example, we documented that an intact axoneme (~200 nm in max height) loses most of its tubules upon MCAK addition (SI Appendix, Fig. S9G). Occasionally, we were able to capture intermediates in this reaction due to adsorption of the dissociated microtubules on mica (Fig. 5F, SI Appendix, Fig. S9H, Video 7). In the example shown in Fig. 5F-G, MCAK addition resulted in the rapid unfurling of the axoneme and the scattering of microtubules on the surface.

These data suggest that the preferential and rapid depolymerization of one tubule in a doublet by enzymes such as MCAK may result in disintegration of the structure by breaking the links between doublets in an axoneme.

## Discussion

Collectively, this study demonstrates the power of AFM imaging in visualizing dynamic processes within dense microtubule arrays in real time and at spatial resolution that, for the first time, allowed observations of individual protofilaments. AFM imaging of depolymerizing microtubule arrays revealed previously unseen structural dynamics and provided new mechanistic insights into how differences in the activities of microtubule remodeling factors at the level of protofilaments and microtubules can lead to differential fate of complex multi-microtubule arrays.

Our data show that the prototypical members of two major depolymerases of the kinesin superfamily, the kinesin-13 MCAK and the kinesin-8 Kip3p, depolymerize complex microtubule arrays in distinct ways. How is this achieved at the mechanistic level? The answer to this question has not been clear despite greater than two decades of enzymatic and structural studies. From the perspectives of enzymology and substrate preference, Kip3 and MCAK are similar. First, both enzymes use family specific loops in the motor domains to target curved microtubules ends (26, 31). Second, depolymerization catalyzed by both motors is not coupled to ATP hydrolysis. Instead, ATPase is inhibited at the microtubule end, and the role of hydrolysis is primarily to dissociate the enzymes from tubulin dimers after depolymerization, recycling for the next depolymerization reaction (26, 51). While an obvious difference lies in the diffusive scanning of MCAK and the processive motility of Kip3 on microtubules, these differences reflect the mechanism by which the enzymes arrive at microtubule ends and not their depolymerase activity (22–25). Indeed, it is observed that a non-motile Kip3 can depolymerize microtubules (26). Our data suggest that the observed differences stem from distinct substrate specificity of the depolymerases at the protofilament level (Fig. 6). We propose that MCAK molecules can effectively depolymerize an exposed protofilament end segment that it encounters at the middle or end of the lattice. This is reflected in the observation that MCAK can propagate defects and protofilaments can be asynchronously depolymerized from the ends. In contrast, a short exposed protofilament end like in a defect, is not sufficient for Kip3-mediated depolymerization from that site. Instead, Kip3 molecules synchronously depolymerize from the multi-protofilament ends of microtubules, where they accumulate. This is consistent with the previously observed co-operativity in Kip3 activity (52, 53). This difference in protofilament-level substrate preference of the two enzymes lead to distinct action of crosslinked microtubules and doublets despite similarities in tubulin binding and ATPase activity at microtubule ends.

**Fig. 6.**
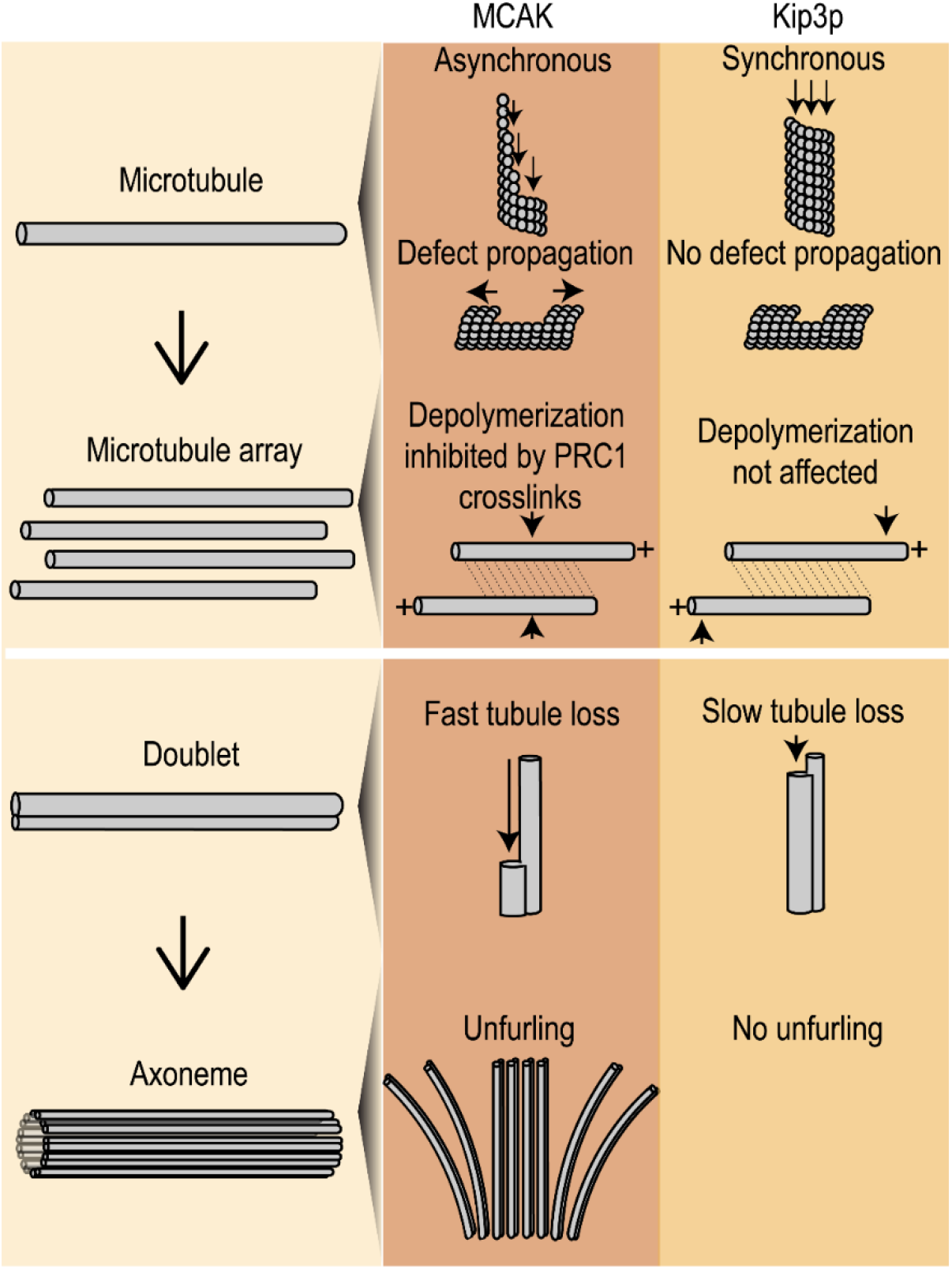
Summary of AFM findings. Schematic of AFM imaging illustrates the distinct structural dynamics and intermediates during microtubule depolymerization with different enzymes and its impact on microtubule arrays. In the context of an individual microtubule: (1) the depolymerization of protofilaments is asynchronous with MCAK and synchronous with Kip3p, and (2) MCAK propagates lattice defects whereas Kip3p does not. In the context of microtubule arrays: (1) Crosslinking by PRC1 protects microtubules against depolymerization by MCAK and does not significantly influence the depolymerization by Kip3p, and (2) MCAK depolymerizes doublet microtubules and results in the destabilization of axonemal arrays. This arises from fast depolymerization of one tubule which compromises the stability of the cylindrical doublet array.

Our results also provide insights into the structural mechanism by which these depolymerases differentially regulate “microtubule aging”. Aging refers to the observation that catastrophe results from a multi-step process, and its probability increases with microtubule lifetime (29). MCAK reduces this process to a first order reaction while Kip3 simply accelerates the rate of steps leading to catastrophe, and consequently MCAK is able to disassemble arrays faster than Kip3 (29). Current models propose that the catastrophe vulnerable state arises from the accumulation of defects and flayed protofilament ends that have fewer laterally stabilized protofilaments (40, 54, 55). How might MCAK and Kip3p differently affect lateral stabilization? The protofilament-level substrate specificity and efficient depolymerization of exposed protofilament ends by MCAK would prevent or greatly reduce the inherent lateral stabilization of protofilaments, thus effectively eliminating the need for accumulation of laterally un-stabilized protofilaments over time or aging. On the other hand, the stabilization of curved tubulin at microtubule ends by Kip3 will serve to increase the rate of aging but not the multi-step nature of the process. Together these observations provide a structural mechanism that can serve as a framework for understanding the molecular underpinnings of aging process, and it’s regulation by depolymerizing enzymes.

The difference in substrate preference at the protofilament level has multiple implications for structural dynamics and regulation of microtubules and the arrays, and the role depolymerases play in these processes. First, lattice defects, which are sites with only a few exposed protofilament ends, are propagated by MCAK but not by Kip3p, suggesting that microtubule arrays with higher level of lattice defects may be especially sensitive to MCAK activity. This may be advantageous for disassembly of microtubule arrays by MCAK in conjunction with proteins such as microtubule severing enzymes (56, 57). On the other hand, the inability to act on defects would be advantageous for depolymerases like Kip3 and other homologs of Kip3, such as Kif18A, that suppress microtubule dynamics for length control (58). Second, the restriction of Kip3p activity to microtubule ends makes it less sensitive to the crosslinking of microtubules by PRC1 compared to MCAK. This may arise from lower density of PRC1 crosslinks at microtubule ends, the primary site of Kip3p action, either because of crowding of Kip3p at ends or due to disruption of microtubule geometry at ends. This feature of Kip3p would allow it to act as an effective length-regulator even in the context densely crosslinked microtubule bundles. It is noteworthy that despite protection by PRC1 crosslinks, MCAK remains an overall faster remodeler due to rapid kinetics of tubulin removal and ability to access exposed protofilaments at both ends of microtubules and at defects. The properties of MCAK also allow it to enzymatically depolymerize one of the most stable microtubule structures in the cell, the doublet microtubules (46, 49, 59). We find that axonemal doublet microtubules can be depolymerized by MCAK, with B-tubule preferentially depolymerized over the A-tubule. The lower stability of the B-tubule due to fewer MIPs and incomplete tubule structure (48, 60, 61), likely exposes protofilaments that are effectively depolymerized by MCAK. Consequently, MCAK mediates the rapid destabilization of axonemal arrays. We think that the fast depolymerization of one tubule compromises the stability of the axonemal array by breaking the links between doublets. In contrast, proteins like Kip3p, which preferentially depolymerize singlets over doublets can act as length regulators at the distal cilium tip, which is predominantly comprised of singlet microtubules (62).

As demonstrated here, previously unknown structural dynamics of individual microtubules and protofilaments within complex arrays can be clearly visualized in the AFM time lapse images. In the case of the two major family of depolymerases, we find that functional dichotomy in the action of depolymerases, which determines how micron-scale arrays are remodeled, can arise from differences in enzyme activity on the nanometer-sized protofilaments. These differences in the observed structural dynamics enable a greater diversity of microtubule array remodeling outcomes as would be beneficial in different cellular contexts when either large scale reorganization or fine-tuning of the architecture of cellular structures in required.

## METHODS

### Atomic Force Microscope Experiments

Microtubule adsorption on mica is achieved by increasing the multivalent cation concentration of the buffer as described previously (6). To prepare microtubule bundles, taxol-stabilized or GMPCPP microtubules, PRC1 and BRB80 buffer with an additional 5 mM MgCl_2_ were combined in a tube. The microtubule and protein mixture was quickly spun down for a few minutes and ~20 μL of this mixture was deposited on a mica substrate freshly cleaved by scotch tape. Single microtubule, doublet, and axoneme samples were diluted with BRB80 and 5 mM MgCl_2_ and deposited on mica as described above. After a few minutes of incubation, ~10 μL of BRB80 buffer was added to the mica and to the AFM tip before imaging the sample by AFM. All AFM experiments were carried out by tapping mode in liquid with the Asylum Cypher S with a silicon nitride tip (BL-AC40TS, radius: 8 nm; spring constant: 0.09 N/m; Oxford Instruments).

## ACKNOWLEDGEMENTS

We thank S. Jiang for purifying MCAK and R. Ohi and D. Pellman for generously sharing Kip3p plasmids and proteins. R.S. was supported by the Pew Biomedical Foundation, the Smith Foundation and the NIH Director’s New Innovator Award. This work was performed in part at the Center for Nanoscale Systems (CNS), a member of the National Nanotechnology Coordinated Infrastructure Network (NNCI), which is supported by the National Science Foundation under NSF award no. 1541959. CNS is part of Harvard University.

## DATA AVAILABILITY

All data needed to evaluate the conclusions in the paper are present in the paper or the supplementary materials.

## AUTHOR CONTRIBUTIONS

S.S.W. designed and performed the experiments, analyzed data and cowrote paper; M.M. performed TIRF experiments and analyzed data; J.S.T. assisted with AFM experiments; R.S. conceived of and designed the experiments, cowrote the paper and supervised the project. All authors discussed the results and commented on the manuscript.

