## Supplemental Information for "Atomic Force Microscopy Reveals Distinct Protofilament-scale Structural Dynamics in Depolymerizing Microtubule Arrays"

SI Appendix Items for

**This PDF file includes:**

1. Methods and Materials
2. Extended Methods
3. Figures S1-S9
4. Video Legends

† These authors contributed equally to this work.

### 1. Methods and Materials

#### Protein purification

PRC1 was expressed and purified as described previously (1). Fluorescent MCAK plasmid was synthesized and the protein expressed and purified as described (2). Kip3p plasmid and protein were a gift from R. Ohi (University of Michigan). Both proteins were purified using standard Ni-NTA purification protocol followed by Superose-6 size exclusion chromatography.

#### Microtubule polymerization

GMPCPP polymerized and taxol-stabilized microtubules were prepared as described previously (1). Briefly, GMPCPP seeds were prepared from a mixture of unlabeled bovine tubulin, X-rhodamine-tubulin, and biotin tubulin, which were diluted in BRB80 buffer (80 mM PIPES pH 6.8, 1.5 mM $MgCl_2$, 0.5 mM EGTA, pH 6.8) and mixed by tapping lightly. The diluted seeds were transferred to a 37°C heating block and covered with foil to minimize light exposure. After approximately 1 hour, 100 µL of warm BRB80 buffer was added to the microtubules and spun at 75,000 rpm, 10 min and 37°C to remove free unpolymerized tubulin. Following the centrifugation step, the supernatant was discarded, and the pellet was washed by round of centrifugation with 100 µL BRB80. The pellet was resuspended in 16 µL of BRB80 and stored at room temperature covered in foil. 20 µM taxol was added for the preparation of doubly stabilized microtubules. Rhodamine-labeled microtubules were used for both TIRF and AFM experiments.

#### Isolation of axoneme

Sperm flagellar axonemes were isolated and purified from sea urchin Lytechinus pictus (Marinus Scientific, LLC, Newport Beach, CA), according to the procedures of Salmon et al. (3). Briefly, sea urchin sperm were collected by inducing them to spawn by injecting the animal with 0.5 M KCl. The sperm were diluted with artificial sea water and put on ice for 20 mins. The sperm suspensions were then centrifuged at 500 rpm. Afterwards, the supernatant was collected, and the sperm were pelleted by centrifuging at 5000 rpm. The sperm pellets were resuspended by trituration in 20% sucrose to osmotically remove the plasma membranes, and the demembranated sperm were homogenized to break the sperm heads from the tails. The suspensions were centrifuged at 10,000 rpm to pellet whole sperm and sperm heads, and the supernatants recentrifuged at 13,000 rpm to pellet the detached sperm tails. The pellet was stratified into a top white layer that contains the demembranated tails and a bottom yellow layer that contains heads and debris. The white layer was collected and resuspended by trituration in isolation buffer. The resuspended tails break the tails into fragments and were further centrifuged at 10,000 rpm. The top white layer was resuspended by trituration in isolation buffer centrifuged to completely separate the tail fragments from heads and debris until the pellet is a single layer of pure white. The white pellet was resuspended in extraction buffer and homogenized and incubated on ice for 45 min to extract dyneins and central pair MTs from the tail fragments. The extracted axonemes were separated from soluble proteins by centrifuging them at 13,000 rpm. The axoneme pellet was resuspended by trituration in extraction buffer, and the axoneme fragments were re-extracted by incubation. The extracted axonemes were pelleted by centrifugation at 13,000 rpm. The extracted axoneme pellet was resuspended by trituration in isolation buffer containing 50% glycerol. Unless specified, all steps were performed at 4°C, and all centrifugation times were 5-10 min.

Doublet fractionation

Axoneme pellets were taken up in Tris-EDTA-DTT solution (TED, 2 mM Tris, 0.2 mM EDTA and 0.5 mM DTT, pH 7.8). As described in the literature, the TED treatment has shown to remove 40-50% of the protein in the sea urchin doublets, which remain associated in sheets of 9 or fewer (4). The treatment time is empirically determined to obtain a mixture of sheets and isolated doublets.

#### Atomic Force Microscope experiments

##### *AFM imaging*

To acquire static AFM images of the sample, clean regions showing flat microtubule arrays were chosen. To get a high-resolution image of the sample, the scan size was decreased to ~ <1 µm. For time-lapse experiments, after finding a flat microtubule bundle on mica, we zoomed into a 1-2 µm region preferably with a few microtubule ends in the scan area. After acquiring this t=0 image, the scanning was briefly paused, the distance between tip and sample was increased by 100-200 nm and 10-20 µL of the depolymerase was injected on to the surface with a pipette without touching the mica disc to prevent losing the scanned area. The amount of liquid on the mica was monitored to ensure that the sample did not dehydrate, and additional buffer was added to the mica to maintain solution volume during the experiments. A range of enzyme concentrations was tested, and final conditions were empirically determined because some protein is lost to the surface and cantilever in these experiments.

All AFM experiments were carried out by tapping mode in liquid with the Asylum Cypher S with a silicon nitride tip (BL-AC40TS, radius: 8 nm; spring constant: 0.09 N/m; Oxford Instruments). After adding the depolymerase, imaging was started at a frame rate of ~3 mins/frame (256 $\times$ 256 pixels at ~1.5 Hz). In liquid, the drive frequency of the tip was ~ 25 kHz. We kept the scan rate at or below 1.5 Hz and scanned at 50-100 pN tip forces to minimize sample damage. Data were collected on microtubules in various orientations due to random binding to the surface, and the features observed were not specific to microtubule orientation with respect to the scan direction. Image acquisition took place over ~30 mins while constantly maintaining the drive amplitude of the tip throughout the experiment. The drive amplitude was maintained slightly above a point where the tip starts to make contact with the surface. To exclude the possibility that depolymerization is arising from the AFM tip, microtubule bundles or doublets were imaged every 5-10 minutes upon adding depolymerase (SI Appendix Fig. 2D-E, 7G). Because of sample contamination and tip damage, using the same tip for all the experiments is not possible. For each new experiment, a new tip was used, and we ensured that we obtained the same quality of AFM images as determined by the spatial resolution or the width of the sample.

*Data Analysis*

Raw AFM data were processed with the Asylum Research (version 16.14.216) and Gwiddyon software (http://gwyddion.net/). All AFM images (height, phase and amplitude) were flattened and any horizontal scars from scanning artifacts were removed prior to analysis. Only the clearest AFM images were used for analysis. After data processing, the height profiles of the microtubules were obtained from the AFM height image.

The number of neighbors was determined by the number of microtubules which were physically contacting an individual microtubule in parallel in a large bundle. For example, a microtubule with no neighbors nearby has N=0, a microtubule in contact with another microtubule has N=1, and a microtubule in contact with two microtubules has N=2.

The depolymerization rates from AFM images were determined by calculating the average length change along the microtubule between the first frame and the last frame from the height time-lapse images (Fig. 2D-E & 4F, SI Appendix, Fig. S1D-E, S4B, S5C & S8D).

For the rate of defect propagation, after the appearance of a defect, the average length change over time from both edges of a defect was measured. In Fig. 2D, ‘slow’ refers the edge with the smallest change in length relative to the ‘fast’ edge. In Fig. 2E, ‘diameter’ refers to change in length over time around the diameter of the microtubule and ‘length’ refers to the change in length over time along the length of the microtubule.

**In vitro fluorescence microscopy assay**

The microscope slides (Gold Seal Cover Glass, 24 × 60 mm, thickness No.1.5) and coverslips (Gold Seal Cover Glass, 18 × 18 mm, thickness No.1.5) were cleaned and functionalized with biotinylated PEG and non-biotinylated PEG, respectively, to prevent nonspecific surface sticking, according to standard protocols (1). Flow chambers were built by applying three strips of double-sided tape to a slide and attaching the coverslip. Sample chamber volumes were approximately 6–8 μL.

Experiments were performed as described previously (1). Biotinylated GMPCPP microtubules, labeled with rhodamine, were immobilized in a flow chamber by first coating the surface with neutravidin (0.2 mg/ml). To visualize microtubule depolymerization, MCAK or Kip3p and 1 mM ATP were flowed into the chamber in assay buffer (BRB80 buffer supplemented with 5 mM TCEP, 2 mM $MgCl_2$, 0.2 mg/ml k-casein, 4 mg/ml glucose oxidase, 0.35 mg/ml glucose catalase, 1% b-mercaptoethanol, and 5% sucrose), and a time-lapse sequence of images was immediately acquired at a rate of 6-12 frames/min. Data were collected for 5–25 min.

All experiments were performed on Nikon Ti-E inverted microscope with a Ti-ND6-PFS perfect focus system equipped with an APO TIRF 100x oil/1.49 DIC objective (Nikon). The microscope was outfitted with a Nikon-encoded x-y motorized stage and a piezo z-stage, an sCMOS camera (Andor Zyla 4.2), and two-color TIRF imaging optics (Lasers: 488 nm and 561 nm; Filters: Dual Band 488/561 TIRF exciter).

ImageJ (NIH) was used to process the image files. Briefly, raw time-lapse images were converted to tiff files. From these images, individual microtubule depolymerization events were identified and converted to kymographs by the MultipleOverlay and MultipleKymograph plug-ins (J. Reitdorf and A. Seitz; https://www.embl.de/eamnet/html/body_kymograph.html).

The following criteria were used to select microtubules for quantitative analysis: (1) firmly attached to the surface throughout depolymerization; (2) both ends visible in the initial frame; (3) not overlapping with another microtubule or tubulin aggregate. A threshold was applied to each kymograph to distinguish microtubule from background. To calculate depolymerization rates, the derivative of the position versus time coordinates of each external edge of the microtubule in the rhodamine channel was measured using Velocity_Measurement_Tool macro and then converted from pixels/frame to nm/min. Breaks within microtubules were visually identified from kymographs with a triangular area of low (background) intensity within the bright microtubule region, which indicates a break in the microtubule that continues to be depolymerized from both sides. Total microtubule length for each condition was calculated as a sum of the widths of all quantified kymographs, converted from pixels to microns.

#### 2. Extended Methods

**Contents** **Page**


| | | |
|---|---|---|
| (A). | AFM sample preparation | 9 |
| (B). | AFM methodological information | 10 |
| (C). | AFM imaging of a microtubule and protofilament | 11 |
| (D). | AFM image analysis: Depolymerizing microtubule | 12 |
| (E). | AFM image analysis: Microtubule bundle | 13 |
| (F). | AFM image analysis: Depolymerizing microtubule bundle | 14 |
| (G). | References | 15 |

#### AFM sample preparation

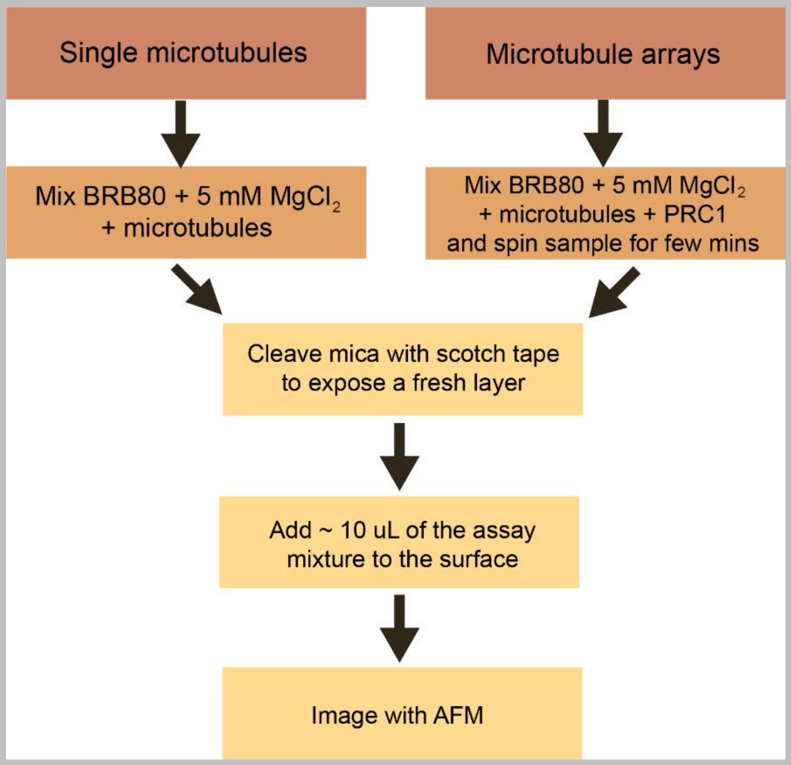


**Extended Data Figure 1. Flow chart of AFM sample preparation for single microtubules and microtubule arrays.**

AFM experiments were carried out on a freshly cleaved mica surface. Mica (polysilicate), a negatively charged surface, is the most widely used substrate for AFM imaging of biological molecules and is used widely to immobilize negatively charged molecules like DNA (5-7).
Microtubules were immobilized non-specifically on the surface in the presence of BRB80 buffer supplemented with 5 mM $MgCl_2$ (Extended Data Fig. 1). No additional treatment of the mica surface was performed. The electrostatic interactions between the $Mg^{2+}$ cations on mica allow for the surface adsorption of negatively charged microtubules. 5 mM $MgCl_2$ is a standard component of microtubule-based assays and does not depolymerize microtubules significantly as observed in our control experiments without enzymes (SI Appendix, Fig. S3).

### AFM methodological information

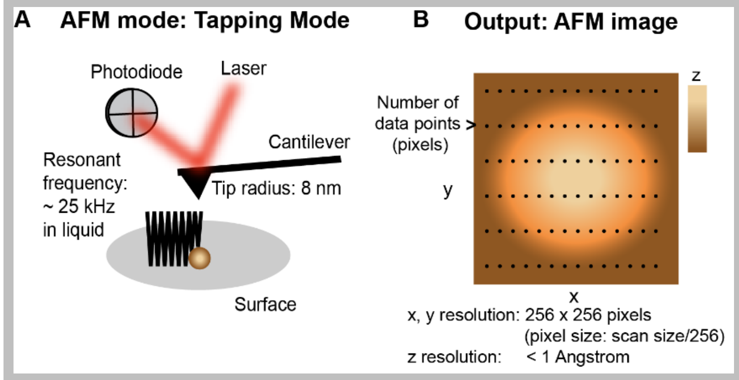


**Extended Data Figure 2. Schematic of AFM experimental set-up, description of the AFM output image and an example AFM image of DNA. A) Schematic of AFM experiment operated in tapping mode. Laser light is reflected off the back of the cantilever to the photodiode. The deflection of the cantilever is measured as it records the surface by gentle tapping of the tip, allowing an image of the topography to be acquired. In liquid, the resonant frequency of the cantilever is 25 kHz and the tip radius is 8 nm. B) Output AFM image of a hypothetical sphere. The AFM image shows the topography in x and y and it is color coded by height (z), by dark to light (brown to cream).**

Tapping mode AFM was used for imaging microtubules. In this mode, which is a gentle technique for imaging soft materials, the cantilever is oscillated near its resonant frequency and makes intermittent contact with the surface while scanning (Extended Data Fig. 2A). The AFM image which is color scaled with height is determined by the Z piezo movement required to maintain the setpoint amplitude interactions. When the probe tip experiences changes in topography, the amplitude feedback changes the Z piezo height to re-establish the amplitude setpoint. The resolution of the AFM image is described by the lateral resolution (x,y) and vertical resolution (z) (Extended Data Fig. 2B). As discussed below, we exploit the high z-resolution for distinguishing microtubules and protofilaments in bundles.

There are two features that determine the x,y resolution of the AFM image: tip shape and pixel size.

1. Tip shape: The radius of curvature and the aspect ratio (radius/length) determine the highest lateral resolution obtainable with a specific tip. The smaller the radius of curvature, the smaller the feature that can be resolved. In our experiments, we use a tip of 8 nm radius.
2. Pixel size: In the AFM output image, the x,y resolution is determined by pixel size in that features smaller than the pixel size of the image cannot be resolved (Extended Data Fig. 2B). In our experiments, the x, y resolution in our images is 256 x 256 pixels, where the pixel size is determined by the scan size divided by 256. For instance an AFM image with a 1 x 1 µm scan size (x,y) is 1 µm/256 pixels = ~4 nm/pixel. In our experiments, the pixel size is ~ 3-4 nm for the high-resolution imaging of microtubules, doublets and axonemes, and ~10-20 nm for the microtubule bundles and axonemes.

The z resolution is determined by the resolution of the vertical piezo scanner movement which has less than 1 Å sensitivity with a baseline system noise <40 pm. We take advantage of the cylindrical shape of the microtubule and the high resolution in z compared to the x-y to differentiate between, singles, doubles, or components/filaments of tubules. This is also advantageous because the lateral x-y resolution and measurements are impacted to some degree by mobility of the tubules, particularly in the scan directions, dilating lateral measurement values. The loss of protofilaments results in the change in height along the short axis of the microtubule, leading to change in the pattern of the topography along this axis. We use this unique feature of AFM to resolve protofilaments within microtubules. The same aspect of AFM imaging allows us to reliably image microtubules within larger arrays by monitoring the height along the array. We illustrate this with examples in the next section.

#### AFM imaging of a microtubule and protofilament

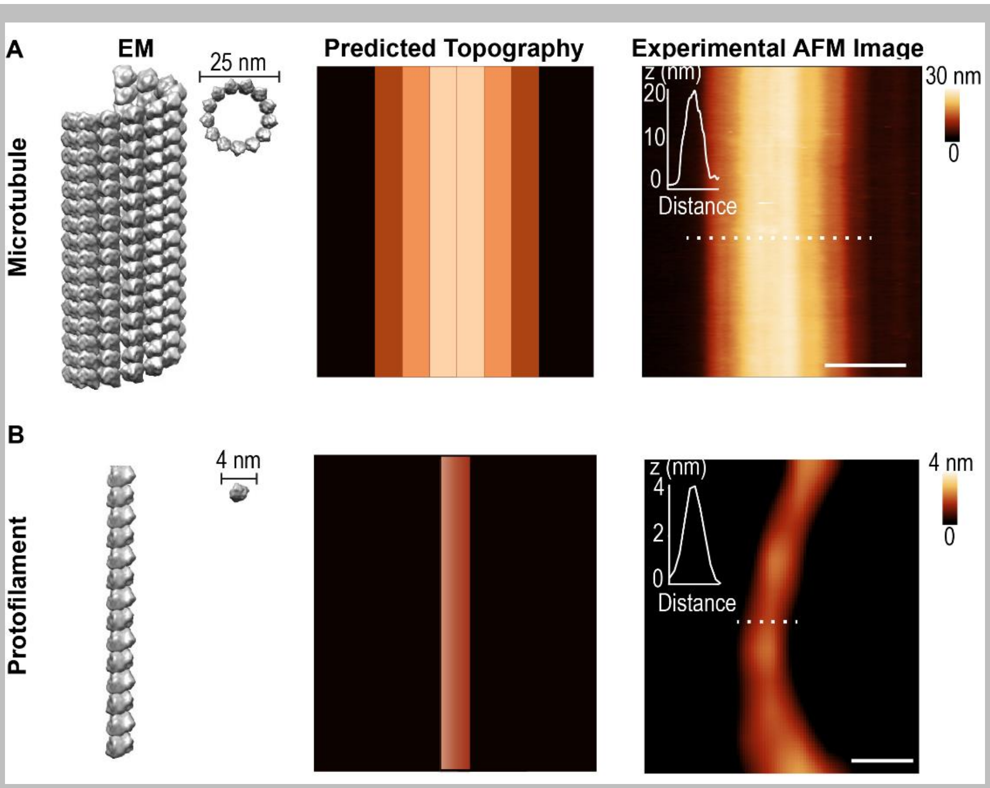


**Extended Data Figure 3. EM structure, estimated topography and high-resolution AFM image of a A) microtubule and B) protofilament. The height profiles in the inset of the AFM images were drawn along the dotted line. The x-y scale bar in A) is 100 nm and B) is 20 nm. The z-scale bar in A) is 30 nm and B) 4 nm.**

Before imaging by AFM, we can estimate the topography of a single microtubule or protofilament from EM images. Based on EM structure of a microtubule, the topography of the microtubule will be characterized by 1) it's cross-sectional height (~25 nm) and 2) striations from protofilaments depending on the scan size. Extended Data Fig. 3A shows EM images of a microtubule and the corresponding estimated topography. As expected, the maximum cross-sectional height of microtubules is ~25 nm at all scan sizes used in this work. At 1 μm scan size, striations along the length of the microtubule, corresponding to protofilaments, are evident. Similarly, based on the EM structure, we expect the cross-sectional height of a protofilament to be 4 nm. As shown in Extended Data Fig. 3B, this is in agreement with our experimental observation.

#### AFM image analysis: Depolymerizing microtubule

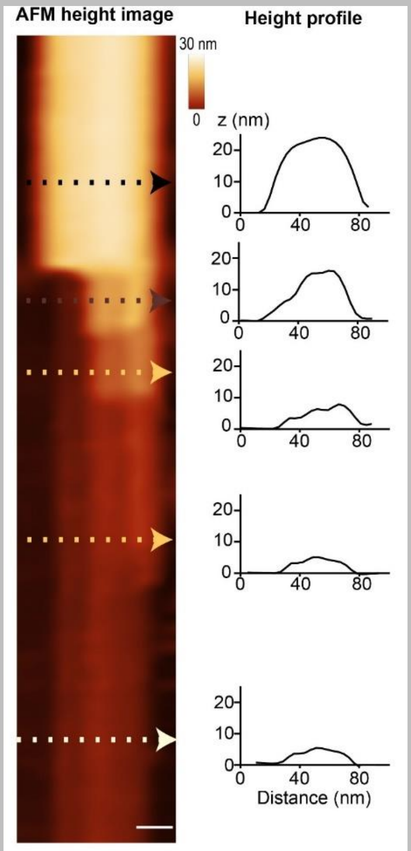


**Extended Data Figure 4. AFM imaging of a depolymerizing microtubule.**

**High resolution AFM image of a depolymerizing microtubule and its corresponding height profiles along the indicated lines on the different sections of the microtubule. Peaks with heights less than 20 nm indicate protofilaments. Scale bar is 20 nm. The x-y scale bar is 30 nm. The z scale bar is 30 nm.**

Since the z resolution of AFM is sub-nanometers, we can discern structures on the order of 4 nm readily even when the lateral resolution is lower in the images at larger scan sizes. This feature is exploited in our experiment to observe loss of protofilaments in microtubules within larger arrays where we need to image a larger field of view. This is illustrated in Extended Data Fig. 4 where the partially depolymerized section of a single microtubule has segments where the topography corresponds to 1-2 or more protofilaments and these can be distinguished based on the topographical height changes. To summarize, the change in height along the z-axis due to microtubule depolymerization allows us to observe the depolymerization of protofilaments by measuring changes in the surface topography over time.

#### AFM image analysis: Microtubule bundles

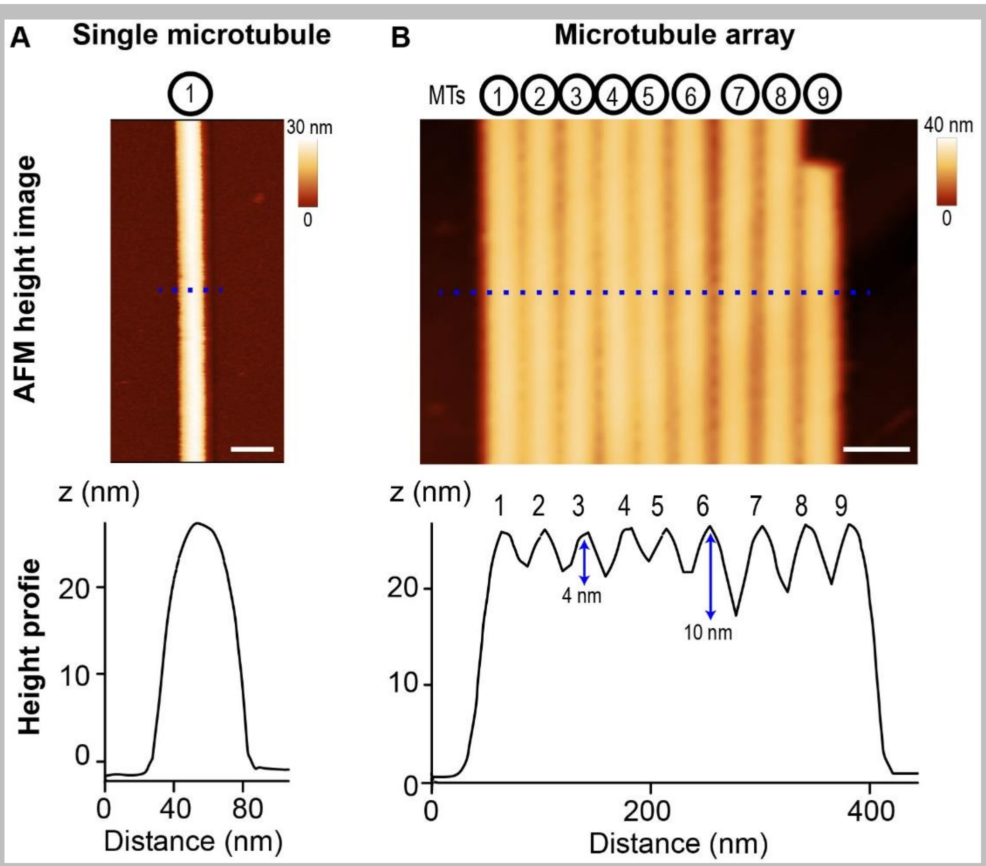


**Extended Data Figure 5. AFM imaging of microtubule bundles. Comparison of A) single microtubule and B) microtubule bundle and its corresponding height profiles. The x-y scale bar in A) is 40 nm and B) is 60 nm. The z scale bar in A) is 30 nm and B) is 40 nm.**

Imaging bundles of microtubules necessitates imaging a larger field of view. While this reduces the lateral resolution (nm/pixel), the spacing between microtubules in the array results in a change in height between neighboring microtubules which is much greater than the z-axis resolution of AFM. As shown in Extended Data Fig. 5, each microtubule in an array with nine microtubules can be clearly identified (numbers). The spacing between the microtubules determines the change in height. In a microtubule array, the crosslinking distance sets the distance between neighboring microtubules in the bundle, and therefore specifies the change in height while moving from one microtubule to the next. In the case of PRC1, the crosslinking distance [~35 nm; (1)] is larger than the diameter of a microtubule. This results in a change in height of 4 nm or more, which readily allows us to distinguish neighboring microtubules in a bundle.

#### AFM image analysis: Depolymerizing microtubule bundle

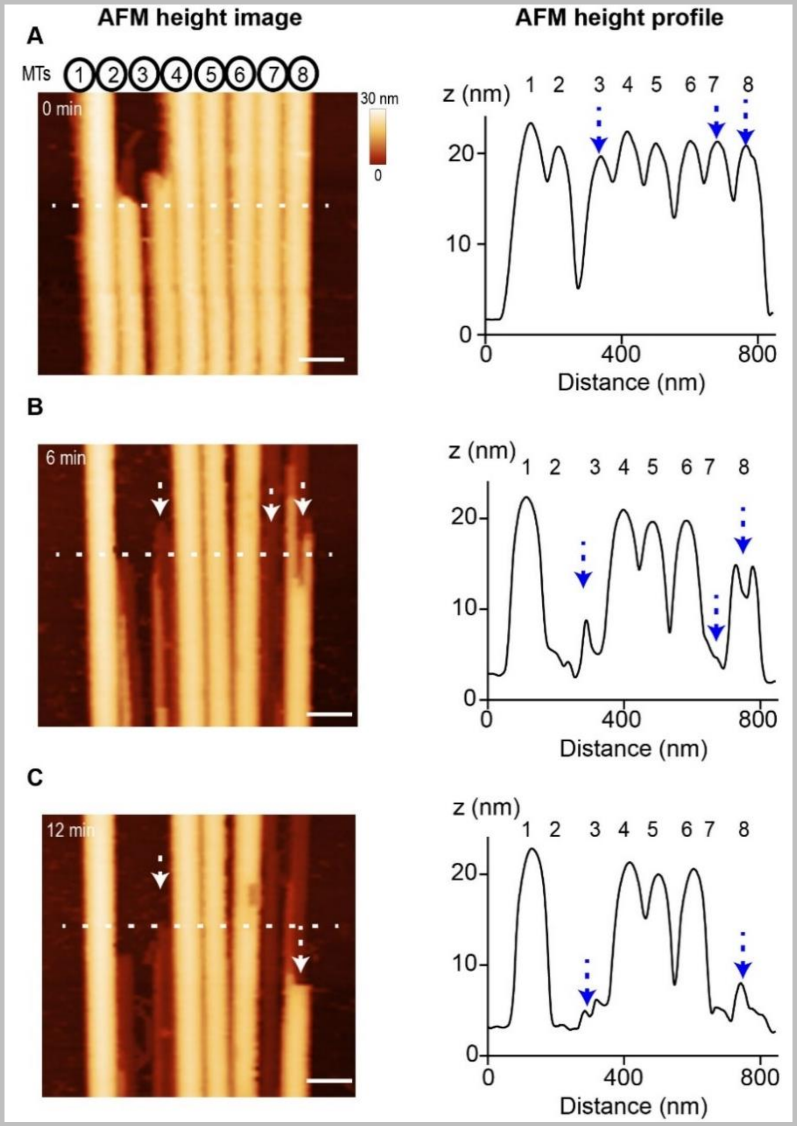


**Extended Data Figure 6. AFM imaging of depolymerizing microtubule array. AFM images of a depolymerizing microtubule bundle at A) 0 min B) 6 min and C) 12 min and its corresponding height profile along the indicated dotted line. The blue arrows in the plots indicate the individual depolymerizing microtubules and the change in height within the bundle over time. Peaks of heights less than 20 nm indicate protofilaments. The x-y scale bar is 50 nm. The z scale bar in is 30 nm.**

When proteins such as PRC1 are bound to microtubules, it can cause changes in the surface topography corresponding to the molecular dimensions of the protein. This along with the larger field of view needed to image larger arrays makes it a challenge to image individual protofilaments within a single intact microtubule. However, loss of protofilaments in a depolymerizing microtubule is readily visualized. This is because the change in topography with loss of protofilaments due to the change in height is significantly greater than the z resolution of AFM imaging. This allows us to reliably image depolymerizing protofilaments within PRC1-bound microtubules. This is illustrated in Extended Data Fig. 6.

### 3. SI Appendix Figures

**Fig. S1**

TIRF

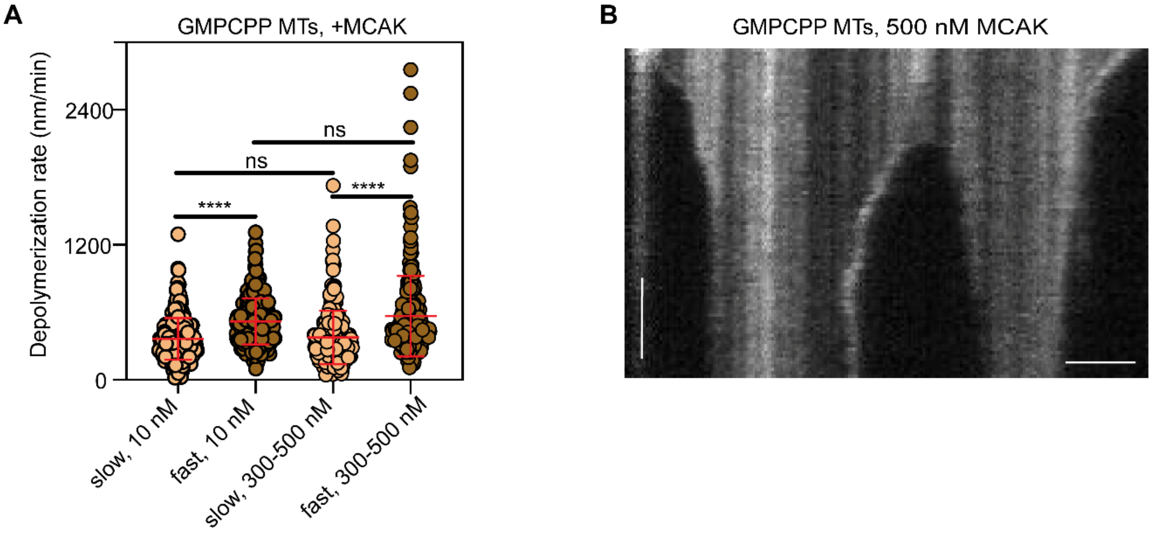


C

| Depolymerase | Concentration (nM) | Number of microtubules | Slow end depolymerization rate (nm/min) | | Fast end depolymerization rate (nm/min) | | Microtubule breaks per mm |
|---|---|---|---|---|---|---|---|
| MCAK | 10 | 281 | 368 | ±95 | 515 | ±113 | 2.97 |
| | 300 to 500 | 252 | 384 | ±91 | 574 | ±203 | 14.30 |

AFM

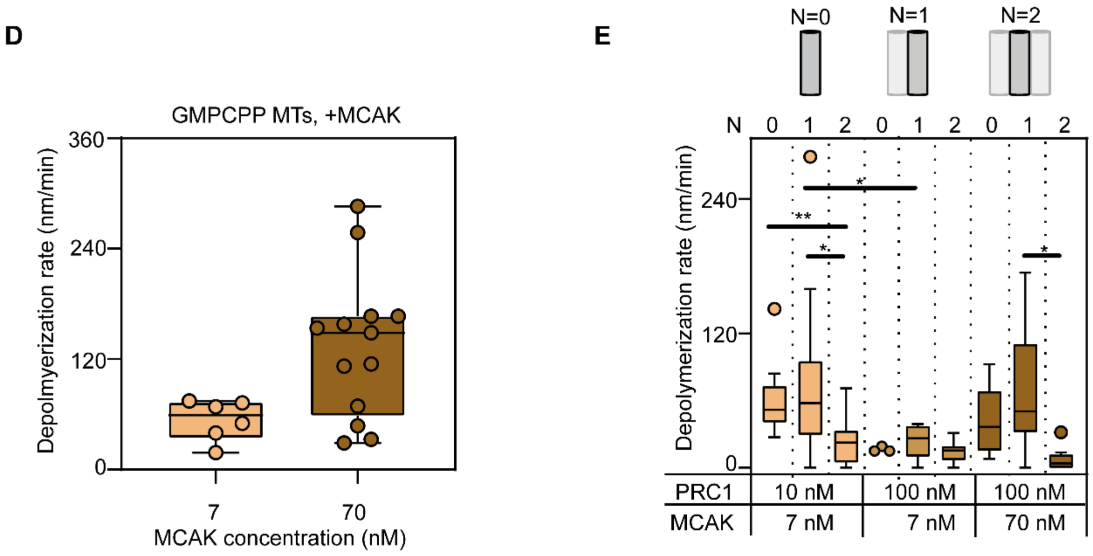

**Fig. S1. Depolymerization of GMPCPP microtubule bundles with MCAK. Related to Figures 1 and 2.**

(A). Depolymerization rates for both ends of microtubules with low or high concentrations of depolymerase MCAK from TIRF experiments. With 10 nM MCAK (n=281 microtubules), the presumed slow-end depolymerization rate is 368±95 nm/min, while the presumed fast-end depolymerization rate is 515±113 nm/min. With high concentrations, ranging from 300 nM to 500 nM, of MCAK (n=252 microtubules), the presumed slow-end depolymerization rate is 384±91 nm/min, while the presumed fast-end depolymerization rate is 574±203 nm/min. Statistical calculations used an unpaired t-test with Kolmogorov-Smirnov correction for non-Gaussian distribution. **** indicates a P-value of < 0.0001. ns indicates a P-value > 0.05. ns indicates a P-value > 0.05.

(B). Representative kymograph of rhodamine-labeled microtubules in the presence of 500 nM MCAK from TIRF experiments. The x-scale bar is 2 µm and the y-scale 2 mins.

(C). Table of slow and fast end depolymerization rates and the number of microtubule breaks per mm with MCAK measured with TIRF.

(D). Box plots of the depolymerization rates of single microtubules from AFM experiments (Median: GMPCPP microtubules, MCAK: 7 nM: 54 nm/min, n=6; 70 nM: 133 nm/min, n=13).

(E). Box plots of the depolymerization rates of neighbor protection analysis in the presence of different PRC1 and MCAK concentrations for neighbors *N*=0, 1, 2 (Median: GMPCPP: PRC1: 10 nM; MCAK: 7 nM: *N*=0: 61 nm/min, n=9; *N*=1: 74 nm/min, n=19; *N*=2: 24 nm/min, n=13; PRC1: 100 nM; MCAK: 7 nM: *N*=0: ~16 nm/min, n=3; *N*=1: 24 nm/min, n=5; *N*=2: 14 nm/min, n=8; PRC1: 100 nM; MCAK: 70 nM: *N*=0: 41 nm/min, n=5; *N*=1: 67 nm/min, n=11; *N*=2: 8 nm/min, n=9).

For D and E, the box plots show the median, the inner quartiles, and maximum and minimum values. Statistical calculations used an unpaired t-test with Kolmogorov-Smirnov correction for non-Gaussian distribution. * indicates a P-value of <0.05. ** indicates a P-value of <0.01.

**Fig. S2**

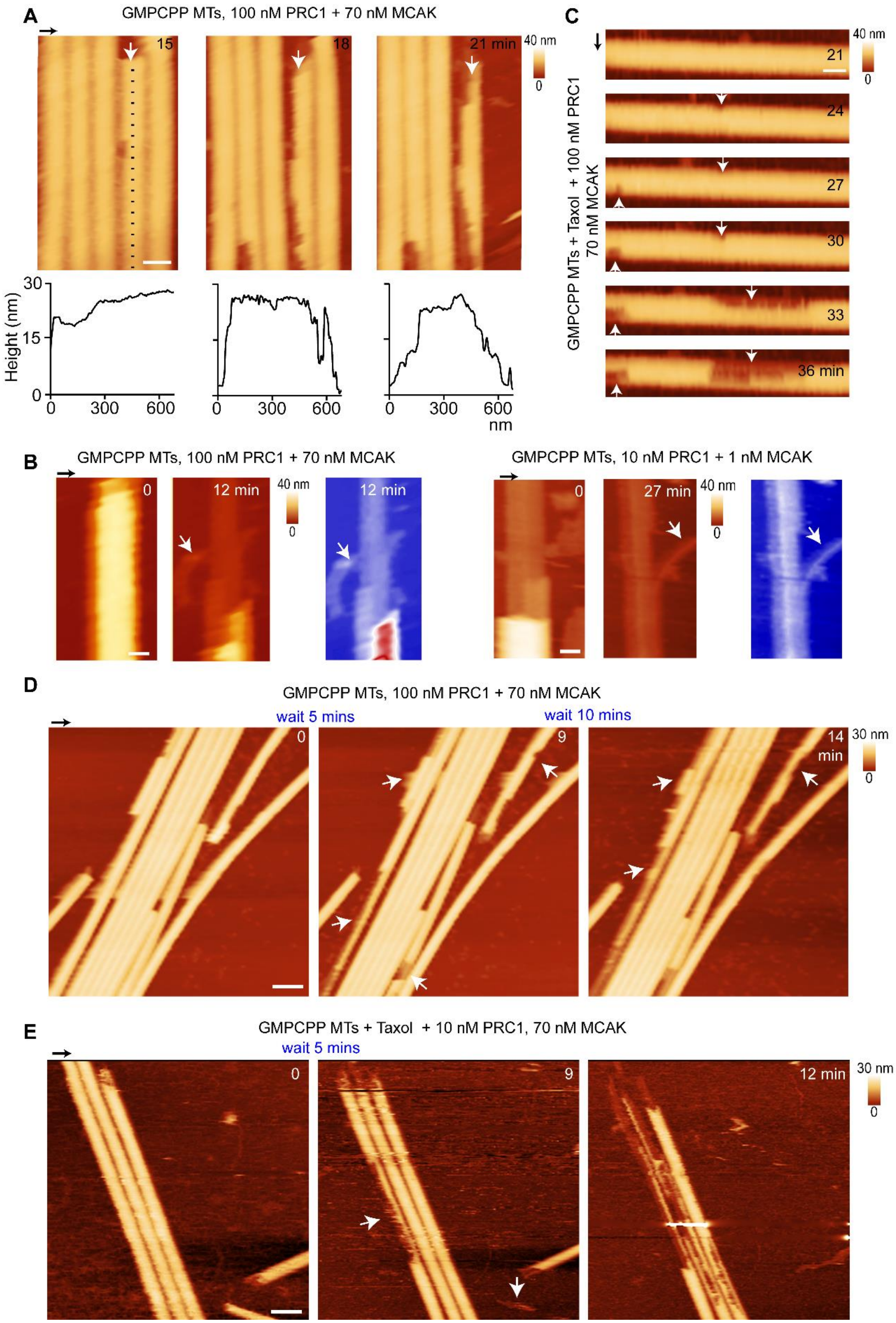

A
GMPCPP MTs, 100 nM PRC1 + 70 nM MCAK
15
18
21 min
40 nm
0
Height (nm)
30
15
0
300
600
nm
C
GMPCPP MTs + Taxol + 100 nM PRC1
70 nM MCAK
21
24
27
30
33
36 min
B
GMPCPP MTs, 100 nM PRC1 + 70 nM MCAK
12 min
GMPCPP MTs, 10 nM PRC1 + 1 nM MCAK
27 min
D
GMPCPP MTs, 100 nM PRC1 + 70 nM MCAK
wait 5 mins
wait 10 mins
9
14
min
30 nm
E
GMPCPP MTs + Taxol + 10 nM PRC1, 70 nM MCAK
wait 5 mins
12 min

**Fig. S2. Depolymerization of GMPCPP and GMCPP + Taxol microtubule bundles with MCAK. Related to Figures 1 and 2.**

(A). Successive AFM images showing depolymerization of individual microtubules within a microtubule bundle by MCAK at the indicated times (GMPCPP microtubules, PRC1:100 nM; MCAK: 70 nM). The x-y scale bar is 40 nm. The corresponding height profiles from the dotted line shows that the stripiness of protofilaments at different heights from the surface.

(B). Successive AFM images showing defect propagation of a microtubule within a bundle by MCAK at the indicated times from the experiment in Fig. 1A. The x-y scale bar is 40 nm.

(C). Ram's horns structures with MCAK + ATP at the indicated time points. (i. GMPCPP microtubules, PRC1:100 nM; MCAK: 70 nM; ii. GMPCPP microtubules, PRC1:10 nM; MCAK: 1 nM). In addition, the AFM height image in blue and red color scale shows the ram's horn structures. The x-y scale bar is 20 nm.

(D). Successive AFM images showing depolymerization of individual microtubules within a microtubule bundle by MCAK after 5-10 min time lapse before each scan (D: GMPCPP microtubules, PRC1:100 nM; MCAK: 70 nM; E: GMPCPP + Taxol microtubules, PRC1:10 nM; MCAK: 70 nM). The x-y scale bar is 40 nm.

The scanning rate is 3 mins/frame with 256 x 256 pixels. The z-scale is from 0 to 40 nm for A-C and 0 to 30 nm for D-E (dark to light brown). The arrow above each panel indicates the scanning direction in the fast axis.

**Fig. S3**

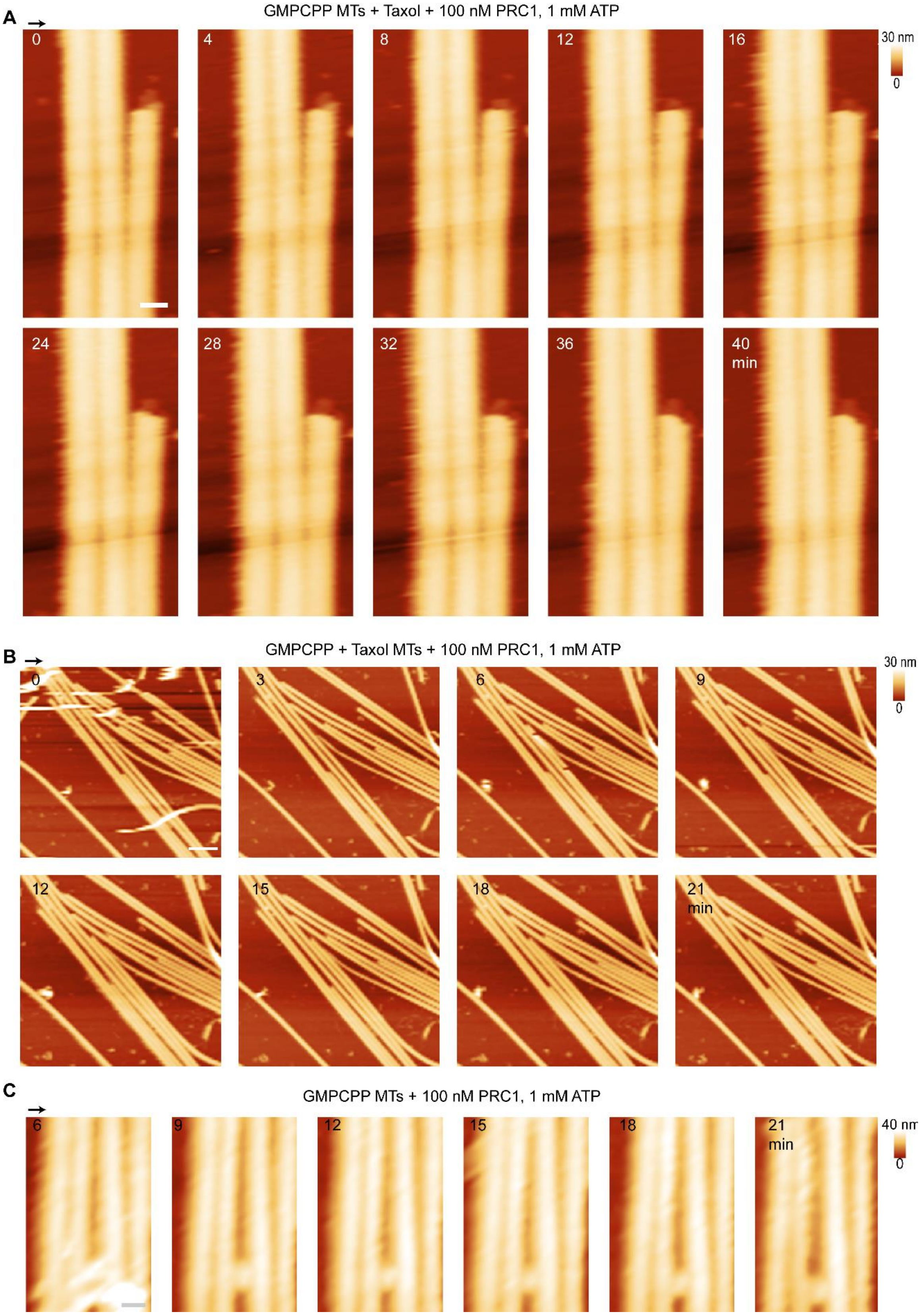
A
GMPCPP MTs + Taxol + 100 nM PRC1, 1 mM ATP
0
4
8
12
16
30 nm
0
24
28
32
36
40
min
B
GMPCPP + Taxol MTs + 100 nM PRC1, 1 mM ATP
0
3
6
9
30 nm
0
12
15
18
21
min
C
GMPCPP MTs + 100 nM PRC1, 1 mM ATP
6
9
12
15
18
21
min
40 nm
0

**Fig. S3. ATP controls of GMPCPP + Taxol and GMPCPP microtubule bundles at different scan sizes. Related to Figures 1, 2 and Figure S2.**

(A). Successive AFM time lapse images showing a PRC1-crosslinked GMPCPP + Taxol microtubule bundle in the presence of ATP alone at the indicated times. The scan size is 1.5 × 1.5 μm. The x-y scale bar is 50 nm. The z-scale is 0 to 30 nm (dark to light brown).

(B). Successive AFM time lapse images showing a PRC1-crosslinked GMPCPP + Taxol microtubule bundle in the presence of ATP alone at the indicated times. The scan size is 4.5 × 4.5 μm. The x-y scale bar is 300 nm.

(C). Successive AFM time lapse images showing a PRC1-crosslinked GMPCPP microtubule bundle in the presence of ATP alone at the indicated times. The scan size is 3 × 3 μm. The x-y scale bar is 50 nm.

The scanning rate is ~3-4 mins/frame with 256 x 256 pixels. The arrow above each panel indicates the scanning direction in the fast axis.

**Fig. S4**

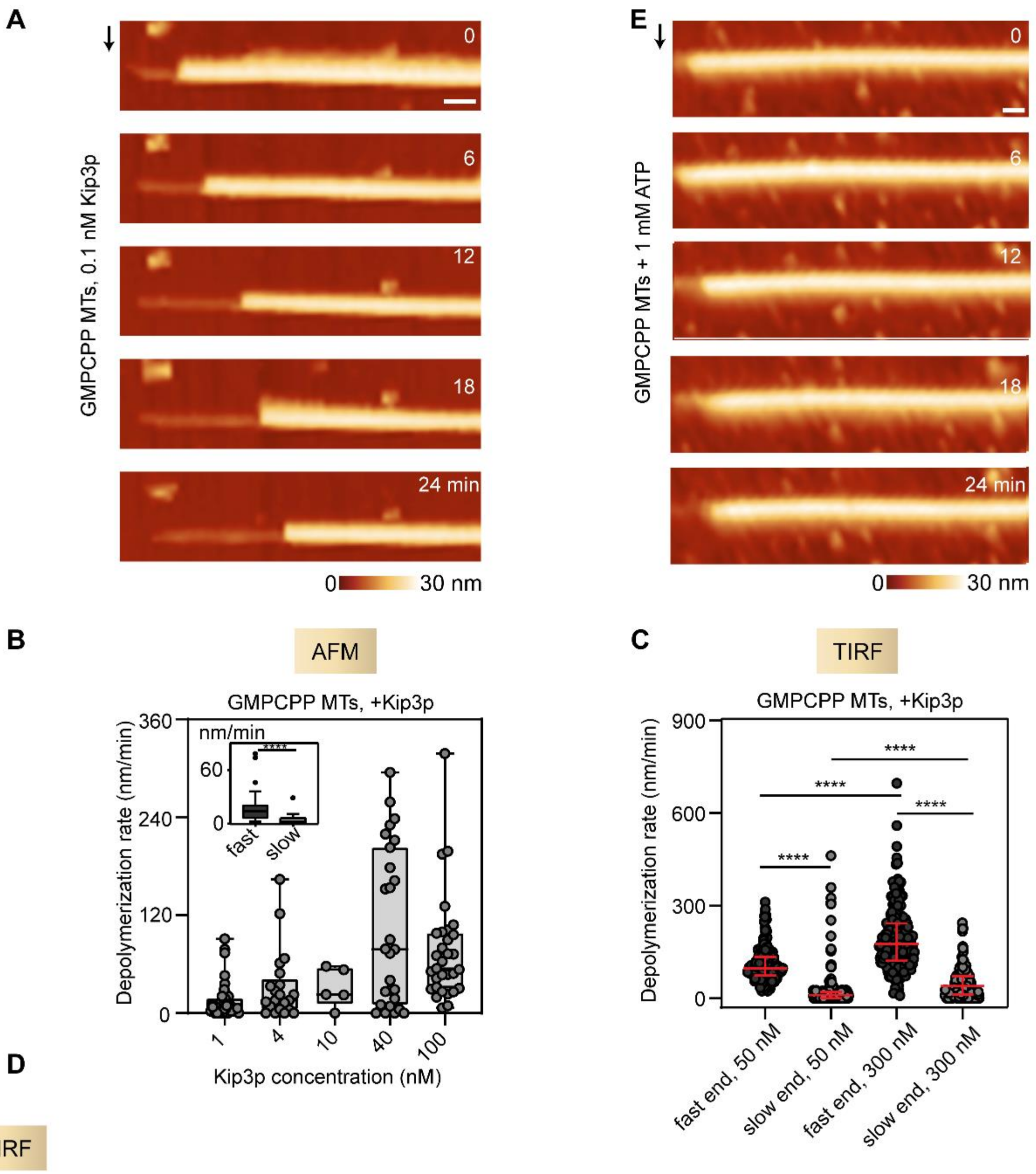


TIRF

| Depolymerase | Concentration (nM) | Number of microtubules | Slow end depolymerization rate (nm/min) | | Fast end depolymerization rate (nm/min) | | Microtubule breaks per mm |
|---|---|---|---|---|---|---|---|
| Kip3p | 50 | 254 | 18 | ±13 | 110 | ±13 | 1.00 |
| | 300 | 177 | 42 | ±14 | 200 | ±33 | 2.34 |

**Fig. S4. Depolymerization of single microtubules with Kip3p. Related to Figure 3.**

(A). Successive AFM images show depolymerization of a single microtubule end by Kip3p at the indicated times (GMPCPP, Kip3p: 0.1 nM). The x-y scale bar is 30 nm.

(B). Depolymerization rates of single GMPCPP microtubules from AFM experiments (Median: GMPCPP microtubules, Kip3p: 1 nM: 11 nm/min, n=35; 4 nM: 16 nm/min, n=17; 10 nM: 31 nm/min, n=5; 40 nM: 103 nm/min, n=27; 100 nM: 58 nm/min, n=28). Inset: The depolymerization rates of 'slow end' and 'fast end' of a microtubule were measured (Kip3p:

1 nM; slow: 5 nm/min, n=23; fast: 20 nm/min, n=23). The box plot shows the median, the inner quartiles, and maximum and minimum values.

(C). Depolymerization rates for both ends of microtubules with low or high concentrations of depolymerase Kip3p from TIRF experiments. With 50 nM Kip3p (n=254 microtubules), the presumed fast-end depolymerization rate is 110±13 nm/min, while the presumed slow-end depolymerization rate is 18±13 nm/min. With 300 nM Kip3p (n=177 microtubules), the presumed fast-end depolymerization rate is 200±33 nm/min, while the presumed slow-end depolymerization rate is 42±14 nm/min.

(D). Table of slow and fast end depolymerization rates and the number of microtubule breaks per mm with Kip3p measured with TIRF.

(E). Successive AFM time lapse images showing a GMPCPP microtubule in the presence of ATP alone at the indicated times. The x-y scale bar is 40 nm.

In panels A and E, the scanning rate is ~3 mins/frame with 256 x 256 pixels. The z-scale is 0 to 30 nm (dark to light brown). The arrow on the left of each panel indicates the scanning direction in the fast axis. For C, statistical calculations used an unpaired t-test with Kolmogorov-Smirnov correction for non-Gaussian distribution. **** indicates a P-value of <0.0001.

**Fig. S5**

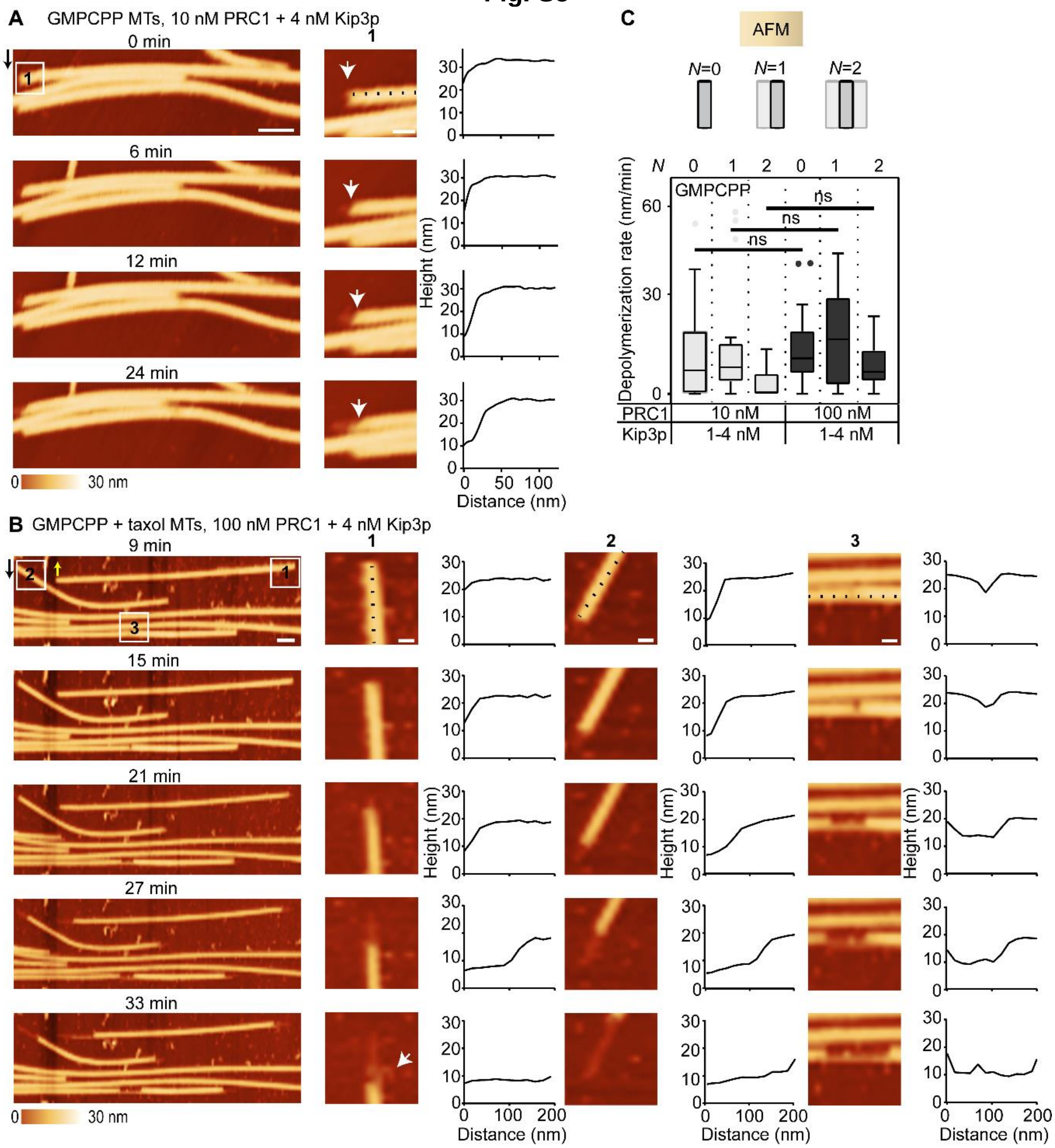


**Fig. S5. Depolymerization of microtubules bundles with Kip3p. Related to Figure 3.**

(A)-(B). Successive AFM time lapse images showing a PRC1-crosslinked microtubule bundle in the presence of Kip3p at the indicated times. Zoomed-in regions of a section of a microtubule end from the experiment shows the absence of stripe-like features at the depolymerizing end (A. box 1; B. boxes 1-3). Protofilament curling (ram's horns) is also observed in B at 33 mins (arrow). The corresponding height profiles from the dotted lines show that the ends of the microtubules and protofilaments were lost synchronously (white arrows) (A. GMPCPP microtubules, PRC1:10 nM; Kip3p: 4 nM; B. GMPCPP + Taxol microtubules, PRC1:100 nM; Kip3p: 4 nM). The x-y scale bar for A-B is 100 nm. The x-y scale bar for zoomed in images A-B is 40 nm.

(C). Depolymerization rates of neighbor protection analysis (see Methods) in the presence of different PRC1 and Kip3p concentrations for neighbors *N*=0, 1, 2 (Median: GMPCPP: PRC1: 10 nM; Kip3p: 1-4 nM: *N*=0: 12 nm/min, n=38; *N*=1: 15 nm/min, n=17; *N*=2: 3 nm/min, n=8; PRC1: 100 nM; Kip3p: 1-4 nM: *N*=0: 13 nm/min, n=25; *N*=1: 17 nm/min, n=17; *N*=2: 8 nm/min, n=14). The box plot shows the median, the inner quartiles, and maximum and minimum values. Statistical calculations used an unpaired t-test with Kolmogorov-Smirnov correction for non-Gaussian distribution. ns indicates a P-value >0.05.

The scanning rate is ~3 mins/frame with 256 x 256 pixels. The z-scale is 0 to 30 nm (dark to light brown). The arrow next to each panel indicates the scanning direction in the fast axis.

**Fig. S6**

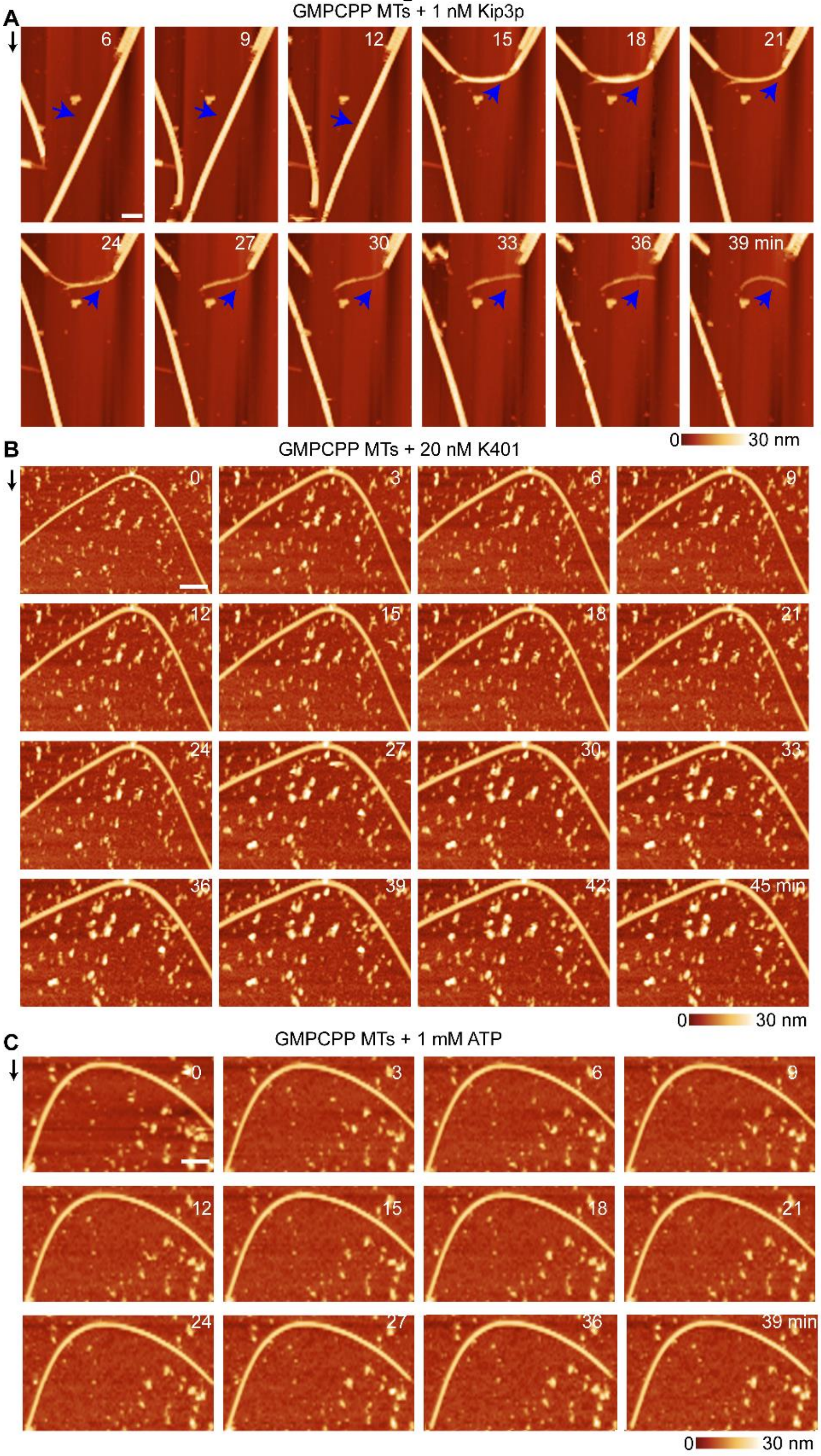
A
GMPCPP MTs + 1 nM Kip3p
6
9
12
15
18
21
24
27
30
33
36
39 min
0
30 nm
B
GMPCPP MTs + 20 nM K401
0
3
6
9
12
15
18
21
24
27
30
33
36
39
42
45 min
0
30 nm
C
GMPCPP MTs + 1 mM ATP
0
3
6
9
12
15
18
21
24
27
36
39 min
0
30 nm

**Fig. S6. Depolymerization of curved microtubules with Kip3p. Related to Figure 3.**

(A). Successive AFM images shows depolymerization of a curved microtubule segment by Kip3p at the indicated times. The blue arrows show a straight microtubule bend into a curve segment at 15 mins, after which undergoes destabilization (GMPCPP microtubules, Kip3p: 1 nM). The x-y scale bar is 80 nm. Related to SI Appendix, Video 4.

(B). Successive AFM images of a curved microtubule segment in the presence of K401 at the indicated times (GMPCPP microtubules, ATP: 1 mM). The x-y scale bar is 300 nm.

(C). Successive AFM images of a curved microtubule segment in the presence of ATP alone at the indicated times (GMPCPP microtubules, ATP: 1 mM). The x-y scale bar is 300 nm.

The scanning rate is ~3 mins/frame with 256 x 256 pixels. The z-scale is 0 to 30 nm (dark to light brown). The arrow next to each panel indicates the scanning direction in the fast axis.

#### Fig. S7

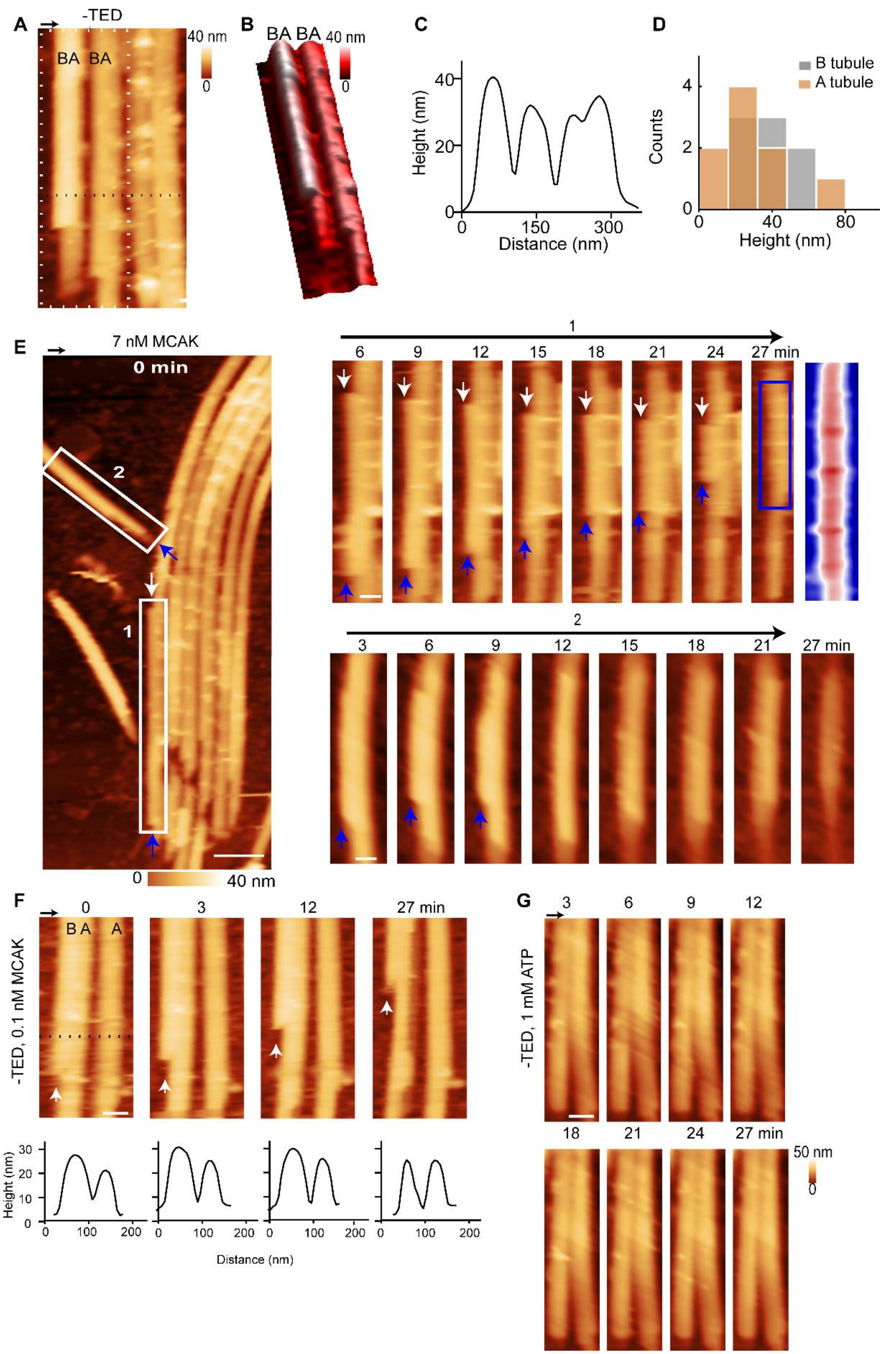
A
-TED
BA BA
40 nm
0
B
BA BA
40 nm
0
C
Height (nm)
Distance (nm)
D
B tubule
A tubule
Counts
Height (nm)
E
7 nM MCAK
0 min
2
1
6 9 12 15 18 21 24 27 min
3 6 9 12 15 18 21 27 min
0
40 nm
F
-TED, 0.1 nM MCAK
0 3 12 27 min
B A A
Height (nm)
Distance (nm)
G
-TED, 1 mM ATP
3 6 9 12
18 21 24 27 min
50 nm
0

**Fig. S7. Depolymerization of microtubule doublets (-TED). Related to Figure 4.**

(A). High-resolution image of a microtubule doublet sheet. ‘A’ and ‘B’ refer to the tubules in the microtubule doublet. The x-y scale bar is 100 nm.
(B). Corresponding 3D image of A (dotted white box).
(C). The corresponding height profile from the dotted line in A.
(D). The height histogram of the ‘A’ and the ‘B’ tubule in a microtubule doublet from AFM images (A=~25 nm; B=40 nm).
(E). An AFM image of a 2D microtubule doublet sheet in the presence of MCAK at t=0 (-TED sample, MCAK: 7 nM). The x-y scale bar is 200 nm. Two examples of bidirectional depolymerization of one tubule in a doublet with MCAK (boxes 1, 2). In 1, the depolymerization of the first end rate= 34 nm/min and second end rate= 13 nm/min of the tubule in the doublet. At 27 min, the magnified area shows that the remaining tubule may be the A tubule due to the periodic features. The x-y scale bar is 80 nm. The z-scale is 0 to 40 nm (blue to red).
(F). Successive AFM height and phase images showing a 2D microtubule doublet and a single tubule in the presence of MCAK at the indicated times. The white arrows show the depolymerization of one tubule (-TED sample, MCAK: 0.1 nM). A and B refer to the tubules in the microtubule doublet. The phase images show periodic features on one tubule which might be outer dynein arms (white arrows). The x-y scale bar is 50 nm. The corresponding height profiles show an asymmetric peak changing into a sharper peak over time.
(G). Successive AFM time lapse images showing a doublet in the presence of ATP alone at the indicated times. The x-y scale bar is 80 nm.

The scanning rate is ~4 mins/frame in A and ~3 mins/frame in E-G with 256 x 256 pixels. In A, E, and F, the z-scale is 0 to 40 nm and in G, 0 to 50 nm (dark to light brown). The arrow above each panel indicates the scanning direction in the fast axis.

### Fig. S8

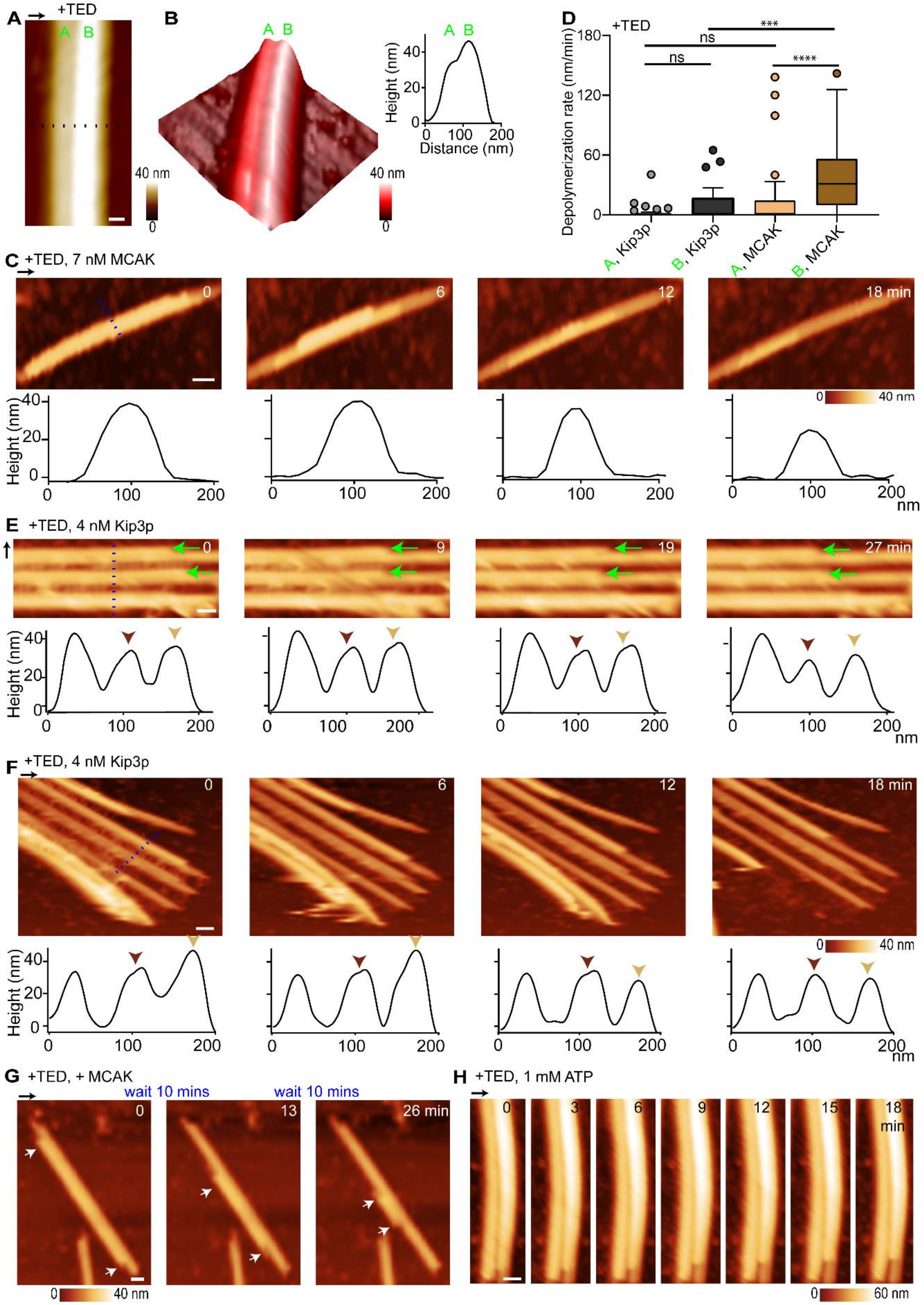

A
+TED
A B
40 nm
0
B
A B
40 nm
0
Height (nm)
Distance (nm)
D
+TED
Depolymerization rate (nm/min)
ns
ns
***
****
A, Kip3p
B, Kip3p
A, MCAK
B, MCAK
C
+TED, 7 nM MCAK
0
6
12
18 min
0
40 nm
Height (nm)
nm
E
+TED, 4 nM Kip3p
0
9
19
27 min
Height (nm)
nm
F
+TED, 4 nM Kip3p
0
6
12
18 min
0
40 nm
Height (nm)
nm
G
+TED, + MCAK
wait 10 mins
wait 10 mins
0
13
26 min
0
40 nm
H
+TED, 1 mM ATP
0
3
6
9
12
15
18 min
0
60 nm

**Fig. S8. Depolymerization of microtubule doublets (+TED). Related to Figures 4 and Figure S7.**

(A). AFM height image of microtubule doublet (+TED sample). The x-y scale bar is 50 nm.

(B). The corresponding 3D representation of A. The height profile from dotted line in A shows that the heights of the 'A' and the 'B' tubule are 35 and 45 nm. No periodic striations were observed.

(C). Successive AFM images showing a microtubule doublet and its corresponding height profiles in the presence of MCAK at the indicated times. The height profiles over time show the change of the doublet height from 40 to 20 nm (+TED sample, MCAK: 7 nM). The x-y scale bar is 100 nm.

(D). Depolymerization rates of 'A' and 'B' tubules in the +TED sample in the presence of MCAK and Kip3p (Median: Kip3p: 0.1-7 nM; A rate= 2.4 nm/min, n=33; B rate= 10 nm/min, n=31; MCAK: 0.1-7 nM: A rate=16 nm/min, n=33; B rate= 38 nm/min, n=30). The box plot shows the median, the inner quartiles, and maximum and minimum values. Statistical calculations used an unpaired t-test with Kolmogorov-Smirnov correction for non-Gaussian distribution. ns indicates a P-value >0.05. *** indicates a P-value of <0.001. **** indicates a P-value of <0.0001.

(E). Successive AFM images showing 2D microtubule doublet sheet in the presence of Kip3p and its corresponding height profiles at the indicated times (+TED sample, Kip3p: 4 nM). The green arrows show the depolymerization of a tubule in each time frame. The brown arrows show the asymmetric peak from a doublet changing into a sharp peak over time. The x-y scale bar is 50 nm.

(F). Successive AFM images showing 2D microtubule doublet sheet in the presence of Kip3p and its corresponding height profiles (from dotted line) at the indicated times. The brown arrows indicate the asymmetric peaks changing to sharp peak indicating the loss of one tubule (+TED sample, Kip3p: 4 nM). The scale bar is 50 nm.

(G). Successive AFM time lapse images showing a microtubule doublet in the presence of ATP alone at the indicated times. The x-y scale bar is 80 nm.

(H). Successive AFM images showing a microtubule doublet and its corresponding height profiles in the presence of MCAK after 5-10 min time lapse before each scan. The height profiles over time show the change of the doublet height from 40 to 20 nm (-TED sample, MCAK: 7 nM). The x-y scale bar is 80 nm.

The scanning rate is ~4 mins/frame for A and ~3 mins/frame for C, E-H with 256 x 256 pixels. The z-scale is 0 to 40 nm (dark to light brown). The arrow above or next to each panel indicates the scanning direction in the fast axis.

#### Fig. S9

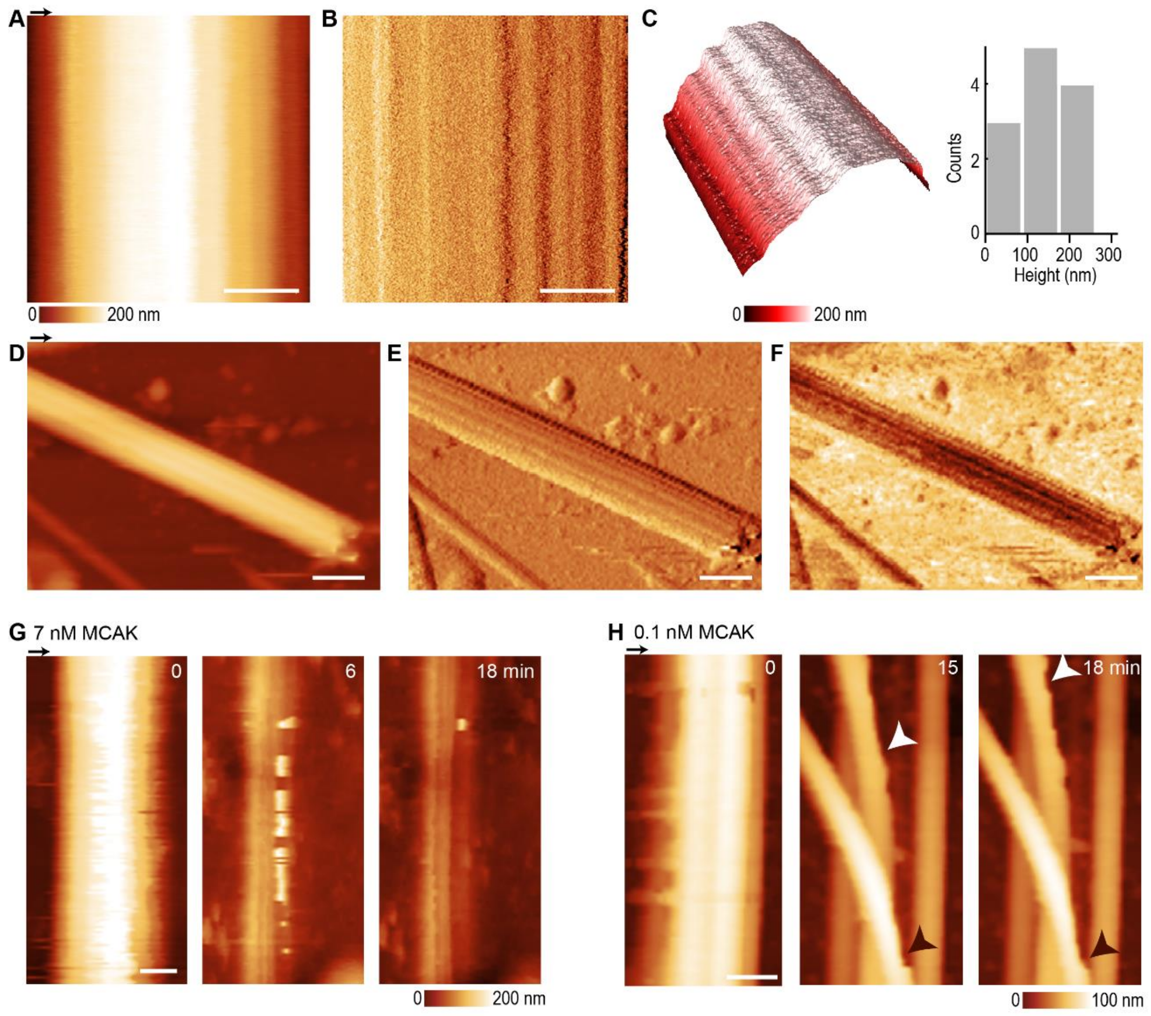


**Fig. S9. Depolymerization of axonemes. Related to Figure 5.**

(A)-(C). Zoomed-in AFM height, amplitude and 3D images of the intact axoneme in Fig. 6A. The height distribution of the axonemes shows a probable peak at around 150 nm. The x-y scale bar is 200 nm. The z-scale is 0 to 200 nm (dark to light brown).

(D)-(F). AFM height, amplitude and phase images of an intact axoneme. This is the initial AFM image before acquiring ATP alone time-lapse experiments. The x-y scale bar is 300 nm. The z-scale is 0 to 300 nm (dark to light brown).

(G). Successive AFM images of an axoneme in the presence of MCAK with 7 nM MCAK at 0-18 mins. These experiments show a rapid loss of tubules. The x-y scale bar is 100 nm. The z-scale is 0 to 200 nm (dark to light brown).

(H). Successive AFM images of a partial axoneme in the presence of MCAK with 0.1 nM MCAK at the indicated times. Zoomed-in regions show the appearance of defects in the tubule (arrows). The x-y scale bar is 150 nm. The z-scale is 0 to 100 nm (dark to light brown).

The scanning rate is 4 mins/frame for A-C and 3 mins/frame for D-H with 256 x 256 pixels. The arrow above each panel indicates the scanning direction in the fast axis.

#### 4. Video Legends

**SI Appendix, Video 1. Depolymerization of a PRC1-crosslinked microtubule bundle by MCAK. Related to Fig. 1E.**
Conditions: GMPCPP + Taxol microtubules, PRC1:100 nM; MCAK: 70 nM: 3 mins/frame. The x-y scale bar is 150 nm. The z-scale is 0–40 nm.

**SI Appendix, Video 2. Depolymerization of a PRC1-crosslinked microtubule bundle by MCAK. Related to Fig. 2A.**
Conditions: GMPCPP + Taxol microtubules, PRC1:100 nM; MCAK: 70 nM: 3 mins/frame. The x-y scale bar is 150 nm. The z-scale is 0–40 nm.

**SI Appendix, Video 3. Depolymerization of a PRC1-crosslinked microtubule bundle by Kip3p. Related to Fig. 3A.**
Conditions: GMPCPP microtubules, PRC1:100 nM; Kip3p: 4 nM: 3 mins/frame. The x-y scale bar is 250 nm. The z-scale is 0–40 nm.

**SI Appendix, Video 4. Destabilization of highly curved microtubule regions by Kip3p. Related to SI Appendix, Fig. S6A.**
Conditions: GMPCPP microtubules; Kip3p: 1 nM: 3 mins/frame. The x-y scale bar is 250 nm. The z-scale is 0–30 nm.

**SI Appendix, Video 5. Depolymerization of doublet microtubules by MCAK. Related to SI Appendix, Fig. S7E.**
Conditions: -TED sample, MCAK: 7 nM: 3 mins/frame. The x-y scale bar is 350 nm. The z-scale is 0–40 nm.

**SI Appendix, Video 6. Depolymerization of doublet microtubules by Kip3p. Related to Fig. 4E.**
Conditions: -TED sample, Kip3p: 5 nM: 3 mins/frame. The x-y scale bar is 300 nm. The z-scale is 0–40 nm.

**SI Appendix, Video 7. Depolymerization of axonemes by MCAK. Related to Fig. 5F.**
Conditions: MCAK: 1 nM: 3 mins/frame. The x-y scale bar is 800 nm. The z-scale is 0–90 nm.
